# Long-term single-cell imaging and simulations of microtubules reveal driving forces for wall pattering during proto-xylem development

**DOI:** 10.1101/2020.02.13.938258

**Authors:** René Schneider, Kris van ’t Klooster, Kelsey Picard, Jasper van der Gucht, Taku Demura, Marcel Janson, Arun Sampathkumar, Eva E. Deinum, Tijs Ketelaar, Staffan Persson

**Author notes:** Shared first authorship (René Schneider, Kris van ’t Klooster). Co-senior and corresponding authors. Correspondence to: Tijs Ketelaar, Laboratory of Cell Biology, Wageningen University, Droevendaalsesteeg 1, 6708 PB, Wageningen, The Netherlands,; Staffan Persson, School of BioSciences, University of Melbourne, Parkville 3010, Victoria, Australia,; Eva E. Deinum, Mathematical and statistical methods (Biometris), Wageningen University, Droevendaalsesteeg 1, 6708 PB, Wageningen, The Netherlands.

## Abstract

Plants are the tallest organisms on Earth; a feature sustained by solute-transporting xylem vessels in the plant vasculature. The xylem vessels are supported by strong cell walls that are assembled in intricate patterns. Cortical microtubules direct wall deposition and need to rapidly re-organize during xylem cell development. We established long-term live-cell imaging of single *Arabidopsis* cells undergoing proto-xylem trans-differentiation, resulting in spiral wall patterns, to investigate the microtubule re-organization. The initial disperse microtubule array rapidly readjusted into well-defined microtubule bands, which required local de-stabilization of individual microtubules in band-interspersing gap regions. Using extensive microtubule simulations, we could recapitulate the process *in silico* and found that local recruitment of microtubule-bound nucleation is critical for pattern formation, which we confirmed *in vivo*. Our simulations further indicated that the initial microtubule alignment impact microtubule band patterning. We confirmed this prediction using *katanin* mutants, which have microtubule organization defects, and uncovered active KATANIN recruitment to the forming microtubule bands. Our combination of quantitative microscopy and modelling outlines a framework towards a comprehensive understanding of microtubule re-organization during wall pattern formation.

## INTRODUCTION

The plant vasculature contains xylem cells that are organised in interconnected tubular networks to enable efficient water distribution to plant organs, and that support plant stature (Myburg et al. 2001). All plant cells are surrounded by primary cell walls, which dictate growth direction. Xylem cells are reinforced by an additional wall layer, referred to as a secondary wall, that is deposited in local thickenings that form highly ordered spatial patterns (Turner et al. 2007). Xylem cells subsequently undergo programmed cell death, which leads to the clearing of their cytoplasmic content and the resulting formation of a hollow tube that provides the water-conducting capacity of vascular plants (Meents et al. 2018).

The major load-bearing component of plant cell walls is cellulose; a β-1,4-linked glucan. Cellulose is synthesised by cellulose synthase (CESA) complexes (CSCs) that span the plasma membrane (Schneider et al. 2016). Nascent cellulose chains coalesce via hydrogen bonds, get entangled in the cell wall and further synthesis thus forces the CSCs to move in the membrane (Diotallevi and Mulder 2007). The CSC delivery to, and locomotion within, the plasma membrane is guided by cortical microtubules that presumably are associated with the plasma membrane (Paradez et al. 2006; Crowell et al. 2009; Gutierrez et al. 2009; Watanabe et al. 2015). Cortical microtubules thus directionally and spatially template the cellulose synthesis machinery during cell wall deposition.

Microtubules undergo dynamic re-organization in response to environmental, developmental and physical cues (Vilches Barro et al. 2019; Lindeboom et al. 2013; Sampathkumar et al. 2014). Spatial control of microtubule arrays can be mediated by small GTPases termed Rho of plants (ROPs) in *Arabidopsis*, which are located in the cytoplasm and at the plasma membrane (Fu et al. 2005; Oda and Fukuda 2012b). For instance, a ROP11-based reaction-diffusion mechanism drives localized secondary wall thickenings during differentiation of meta-xylem cells, which produce pitted wall patterns. This mechanism involves local activation of ROP11 at the plasma membrane and recruitment of microtubule depolymerizers, such as MICROTUBULE DEPLETION DOMAIN (MIDD)1 and KINESIN (KIN)13A (Oda and Fukuda 2012a; Oda and Fukuda 2013; Nagashima et al. 2018). The microtubule-associated proteins IQ-DOMAIN (IQD)13 and IQD14 control the shape of the active ROP11 domains whereas BOUNDARY OF ROP DOMAIN (BDR)1 and WALLIN (WAL) direct actin filaments to the border of the active ROP11 domain (Sugiyama et al. 2017; Sugiyama et al. 2019). In contrast to meta-xylem, the proto-xylem (cells that undergo differentiation when the surrounding tissue still elongates) consists of periodic or spiralling secondary wall bands. Mutations in the genes important for pit patterning in meta-xylem do not cause obvious proto-xylem wall defects, indicating a different regulation for periodic band patterning than for pit formation. To support such band patterning, microtubules undergo a transition from a diffuse array with variable microtubule orientations into a banded array where microtubule orientations are homogeneous (Schneider et al. 2017; Watanabe et al. 2015; Watanabe et al. 2018). Here, cortical microtubules mimicked the secondary wall patterns (Turner et al. 2007) and template secondary wall cellulose deposition (Watanabe et al. 2015; Watanabe et al. 2018; Schneider et al. 2017). Nevertheless, the principles by which the dynamic microtubule network is re-organized during the transition from primary to secondary wall deposition in proto-xylem remain elusive. In addition, the transition period of microtubule array configurations during proto- and meta-xylem formation is not well defined.

Xylem differentiation occurs in a sequential manner along the central axis of roots and shoots, which is buried underneath several cell layers. Detailed proto-xylem formation is therefore difficult to visualize. However, Yamaguchi et al. (2010) generated an elegant system in which proto-xylem formation can be induced in cells that typically do not make secondary walls in *Arabidopsis* (Yamaguchi et al. 2010; Yamaguchi et al. 2011). Here, the proto-xylem inducing “master” regulator VASCULAR NAC-DOMAIN (VND)7 is under control of a chemically inducible glucocorticoid-receptor element and can drive ectopic proto-xylem differentiation upon induction. Recently, Watanabe et al. (2015; 2018), Li et al. (2016a), Schneider et al. (2017) and Li et al. (2016b) used this system to investigate the behaviour of secondary wall cellulose synthesis, to measure the impact of CELLULOSE SYNTHASE INTERACTING (CSI)1 on this process and to assess coordination between transcript and metabolite changes, during proto-xylem formation, respectively. This system is thus of great aid in understanding the transition from primary to secondary wall formation during proto-xylem development.

Here, we established long-term single live-cell imaging of *Arabidopsis* epidermal hypocotyl cells undergoing proto-xylem trans-differentiation. We quantitatively analysed microtubule dynamics parameters over time and used these parameters as input for computer simulations to understand the processes that drive the re-arrangement of the microtubule array during proto-xylem formation.

## RESULTS

### Microtubule array re-organization proceeds non-linearly and rapidly during proto-xylem formation

To study the behaviour of the microtubule array upon induction of proto-xylem formation we used an mCHERRY-TUA5 microtubule reporter line in the VND7-inducible background (Supplementary Fig. S1; Schneider et al. 2017). We classified the trans-differentiation progression into early, mid, and late stages based on microtubule array changes (Watanabe et al. 2015), using anisotropy measurements (Supplementary Fig. S2; Boudaoud et al. 2014). These measurements revealed an increased progression of array anisotropy during early, mid, and late stages of proto-xylem formation. However, this classification of array patterns is based on ‘snapshots’ of the microtubules and thus ignores their dynamic behaviour that drives the re-organization. To assess the microtubule dynamics, we developed an automated image acquisition script and optimized our live-cell imaging procedure by implementing a sample chamber that blocked water evaporation, which allowed us to acquire long-term recordings of single cells. The automated script enabled temporal control of the time lapses. We initially observed that microtubule bands emerged approximately 12 hours after VND7 induction. To capture the complete array rearrangements, we therefore started detailed observations before changes to the microtubule array were visible (typically six to 11 hours after induction). We first recorded single images of microtubules every 30 seconds for five hours in a low-temporal-resolution experiment (Fig. 1a, Supplementary Movie 1). We found that multiple microtubule bands formed simultaneously in the cells. This initial pattern was readjusted during the following two hours (Fig. 1a, yellow arrows). Readjustments involved gradual shifting of microtubule bands (Fig. 1b, arrowheads) that sometimes led to two neighbouring bands merging (Fig. 1b, asterisks). After such readjustments, equally spaced and parallel bands remained at fixed positions throughout the rest of the time course.

**Figure 1.**
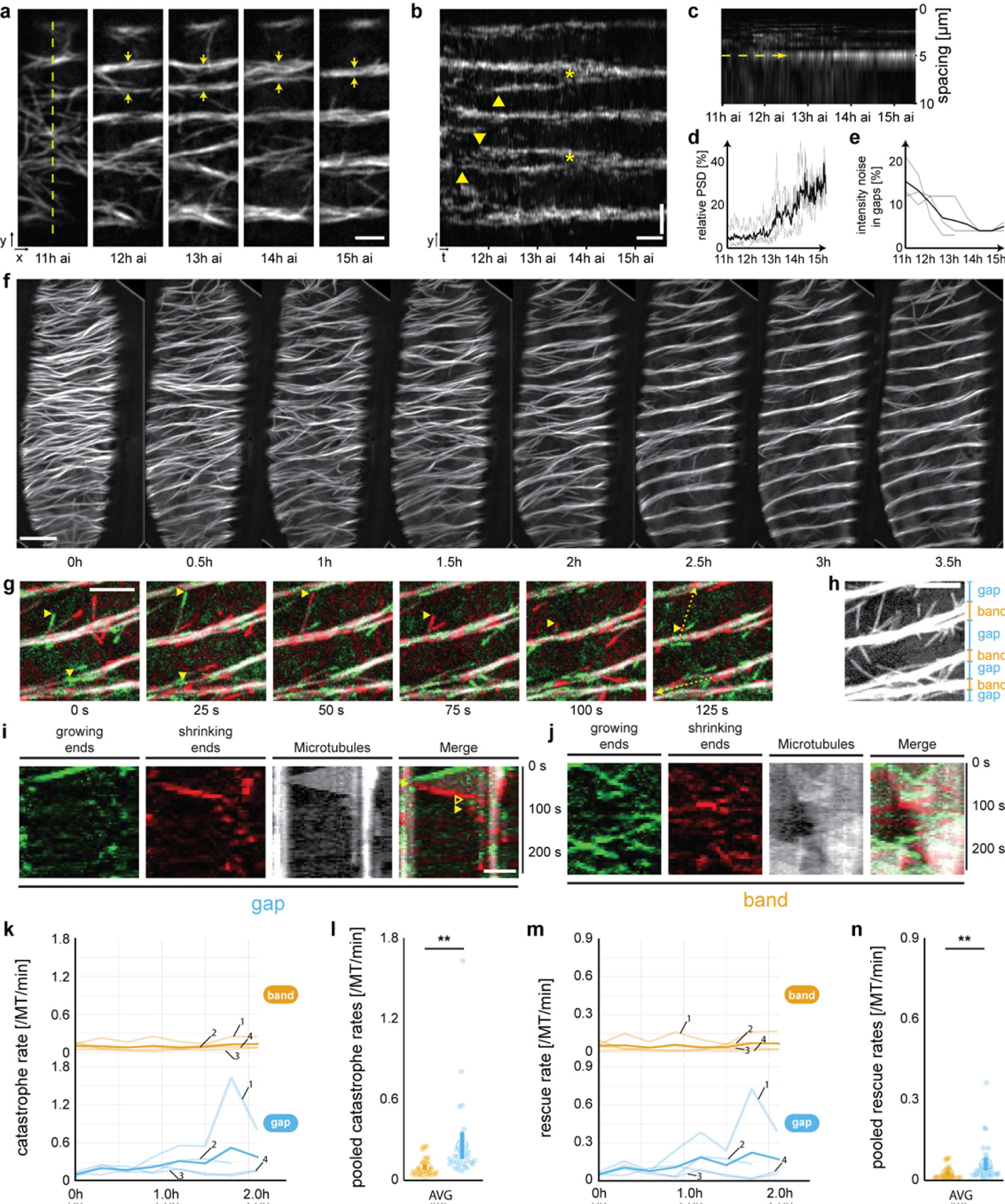
The inducible VND7 system drives ectopic microtubule rearrangements. **a**, Long-term time-lapse recording using spinning disk microscopy of microtubule rearrangements at low temporal resolution (*Δt* = 30 seconds) in an induced hypocotyl cell. Scale bar = 3 µm. Microtubule bands emerge in close proximity to each other but merge into one dominant band over time (anti-parallel pair of arrows). **b**, Kymograph along the dashed line in (a) reveals the emergence (arrow heads) and merging (asterisks) of microtubule bands. Scale bars = 30 min and 3 µm. **c**, Periodogram of the kymograph in (a) reveals development of a dominant microtubule spacing with an average of 5.3 ± 0.5 µm (mean ± s.e.m., 6 cells from 4 seedlings). **d**, The relative power spectral density (PSD) measured at the respective main spacing of four individual cells (grey lines) increases rapidly from 5.5 ± 0.3 % (thick line, mean ± s.e.m.) to 26.9 ± 0.5 % within approximately one hour. **e**, The microtubule noise (standard deviation normalized to sum of intensity of microtubules located in gaps) decreases rapidly in gaps from 15 ± 5 % (thick line, means ± s.d.) to 5 ± 1 %. **f**, Time-averaged images of a long-term time-lapse recording at high temporal resolution (Δ*t* = 5 seconds for 5 minutes every 30 minutes) of YFP-labelled microtubules in an induced hypocotyl cell. Scale bar = 5 µm. **g**, Time-lapse of YFP-labelled microtubules in mid stages of proto-xylem formation. Information on growing (green) and shrinking (red) microtubule ends were extracted using frame difference analysis. Two dynamic microtubule ends are highlighted (arrowheads). Scale bar = 5 µm. **h**, Classification of microtubule bands (orange) and gaps (blue), respectively. **i**-**j**, Kymographs along the dashed lines in (g). In gaps (top kymograph in g), growing microtubule ends undergo catastrophes (filled arrowheads) and become shrinking microtubule ends. Shrinking microtubule ends undergo rescues (empty arrowheads) and become growing microtubule ends. In bands (bottom kymograph in g), microtubule ends predominantly grow anti-parallel along the band axis. Scale bar = 2 µm. **k-n**, Quantification of microtubule dynamics in four independent time series (fine lines, indicated by number in panel k and m) divided into microtubule bands (orange) and gaps (blue): (k, m) individual and (l, n) pooled catastrophe rates *r*_cat_ and rescue rates *r*_res_, respectively. The thickened line represents the time average of all measurements. Data pooled from all 35 time points from four cells (AVG); means ± 95% confidence intervals. Statistics: Welch’s unpaired *t*-test, ** p < 0.005.

To gain insight into the periodicity maintenance during microtubule array patterning, we performed periodogram analysis of the intensity profile along the cell’s growth axis (dashed line in Fig. 1a). A periodogram provides an estimate for how a particular (spatial) frequency contributes to a fluctuating signal. We calculated the power spectral density (PSD) of the intensity profile in Fig. 1a, and plotted it against the spatial frequencies in the profile (Fig. 1c). We found that the PSD initially was evenly distributed, indicating contributions from all spatial frequencies to the intensity profile as expected from a diffuse array. However, the PSD subsequently peaked at spatial frequencies corresponding to approx. 5 µm (dashed arrow in Fig. 1c). By plotting the temporal development of the peak in the PSD, we estimated that the rearrangement of the microtubule array, i.e. from a diffuse array into periodic bands, occurred within one to two hours (Fig. 1d). The microtubule intensity gradually decreased in the gaps between the bands, indicating continuous microtubule removal from the gaps (Fig. 1e). Hence, while the VND7-driven trans-differentiation takes relatively long (up to several days; Supplementary Fig. S3; Kubo et al. 2005; Yamaguchi et al. 2010), the microtubule array re-organization occurs rapidly.

### Microtubule dynamic instability parameters are different between bands and gaps

The low-temporal resolution recordings enabled us to observe global microtubule re-organization during proto-xylem formation. However, individual microtubules undergo dynamics that occur on the timescale of a few seconds and are thus not captured in the long-term recordings. We therefore increased the temporal resolution to 5-second intervals, taken for five minutes every 30 minutes. To avoid photobleaching we generated a YFP-TUA5 expressing VND7-inducible line and used it instead of mCHERRY-TUA5. Similar to above, we found that the total microtubule intensity differed between bands and gaps during the time course (Supplementary Fig. S4). Whereas the intensity remained relatively constant in bands (94 ± 24 % of initial band intensity counts; mean ± s.d. measured throughout whole time course), it decreased within an hour after the start of the recording (48 ± 13 % of initial gap intensity counts) in the gaps and remained low afterwards (37 ± 15 %). The intensity ratio between bands and gaps, which can be interpreted as a measure for band progression, consequently increased throughout the time course.

We were able to record four long-term image series that captured the complete trans-differentiation process (Fig. 1f, Supplementary Fig. S5-S8, and Supplementary Movies 2-5). We found that microtubules underwent frequent depolymerization due to catastrophes when they were located in gaps (Figs. 1g-i). By contrast, the majority of microtubules in the bands appeared to grow, with low depolymerisation frequency (Fig. 1j). To quantify microtubule dynamics in bands and gaps, we used image processing to detect growing and shrinking microtubule ends in the time-lapse recordings (Materials & Methods, Supplementary Fig. S9; Schneider et al. 2019). This approach allowed quantification of microtubule growth (*v*_+_) and shrinkage (*v*_-_) speeds, and catastrophe (*r*_cat_) and rescue (*r*_res_,) rates, which are the most relevant parameters to characterize microtubule dynamics (Tindemans et al. 2014; Tindemans and Deinum 2017). We divided the cortical area into regions that developed into bands and gaps (Fig. 1h) and determined the microtubule dynamic instability parameters (Supplementary Tables 1-4).In bands, microtubules grew and shrank on average with a speed of 53 ± 15 nm/min and −80 ± 19 nm/min, respectively, whereas in gaps, microtubules grew and shrank on average with a speed of 54 ± 15 nm/min and −86 ± 20 nm/min (means ± s.d., 35 time points from four time series). Neither microtubule growth nor shrinkage speeds were significantly different between bands and gaps throughout the microtubule re-organization process (Supplementary Fig. S4). We further measured *r*_cat_ and *r*_res_ and found both to be significantly larger in gaps as compared to bands (Figs. 1k-n). Microtubules in the band regions underwent catastrophes and rescues at average rates of 0.096 ± 0.066 and 0.054 ± 0.048 events per microtubule end per minute, respectively. Significantly higher rates and variations were found in gaps with average *r*_cat_ and *r*_res_ of 0.258 ± 0.282 and 0.120 ± 0.138 events per microtubule-end per minute, respectively (*p*_band_ < 0.002 and *p*_gap_ < 0.011; Welch’s unpaired *t*-test). While we thus observed significant changes in microtubule density and catastrophe/rescue rates in bands versus gaps across the four time series, we also observed variations in microtubule re-organization speed and dynamics between the individual cells (Supplementary Fig. S5-8). Taken together, these results indicate that microtubules display more vigorous dynamics in the gaps as compared to bands.

### A conceptual framework to simulate microtubule band formation

To gain insight into how microtubule dynamics contribute to band formation, we proceeded to establish a framework that could simulate these patterns. To do this, we drew inspiration from three theory-derived concepts (Tindemans et al. 2010): (*i*) the so-called control parameter *G*, which collapses the microtubule dynamic instability parameters and nucleation rate into a single number, (*ii*) the average microtubule length in absence of interactions *L*_0_, and (*iii*) the average microtubule lifetime *τ* (Material & Methods and Supplementary Information). Using the measured microtubule dynamic instability parameters obtained from our time series recordings, and an assumed isotropic nucleation rate, we found that *G, L*_0_, and *τ* were approximately equal between our defined band and gap regions before induction, as expected for a diffuse microtubule array (Supplementary Fig. S5-S8 and S10). However, differences became evident upon microtubule band formation. We estimated *τ* to be 29.4 ± 16.9 minutes in bands (*τ*_band_; means ± s.d., *n* = 35 time points), whereas in the gap *τ* was significantly shorter: 11.0 ± 5.3 minutes (*p* < 0.0001, Supplementary Fig. S10). We used these values to calculate ratios between gaps and bands, *G*_gap_/*G*_band_ and *τ*_band_*/τ*_gap_, to compare the course of the re-organization process in different cells. We found that each cell displayed a phase of peaking ratios but that these peaks occurred at different time points (Supplementary Fig. S5-S8 and S10). The ratios of *G*- and *τ*-values followed very similar curves, reflecting the near inverse relationship between the definitions of *G* and *τ* (Material & Methods and Supplementary Information). Whereas the progression of array behaviour differed between cells, all ratios started close to one, indicating a diffuse microtubule array, and maintained similar overall trends.

### Simulations reveal that arrangements of microtubule nucleation control band patterns

To investigate whether the microtubule-based parameters (*G*, *τ* and *L*_0_) are sufficient to separate a diffuse microtubule array into bands and gaps, and thus to mimic proto-xylem trans-differentiation, we performed computer simulations (Material & Methods and Supplementary Information). To this end, we used an extended version of the ‘*Cortical Sim*’ software (Tindemans et al. 2014) and a parameter set based on the measurements outlined above (Supplementary Tables 1-4). Cortical Sim renders dynamic microtubules as growing and shrinking line segments on a closed surface representing the cell cortex. These “microtubules” interact through frequent collisions, occurring as growing microtubules impinge on obstructing ones. The outcome of such a collision depends on the relative angle: ‘*zippering*’ or ‘*bundling*’ (continued growth along the obstructing microtubule) for angles < 40 degrees and ‘*crossover*’ (continued growth without change of direction) or ‘*induced catastrophes*’ (switch to shrinkage) for larger angles (Dixit and Cyr 2004).

For our simulations, we used 60 µm long and 15 µm wide cylindrical cells (Fig. 2a). Along the long axis, we subdivided the cell into 5 µm wide gaps, consistent with our PSD measurements (Fig. 1c), and 1 µm wide bands, to which we applied different parameters based on the experimentally observed microtubule dynamic instability parameters in the respective regions (Supplementary Table 1-4). To create a transverse and diffuse microtubule array, we implemented a two-hour initiation phase where the microtubule dynamic instability parameters were similar between band and gap regions. Because of the variation among the quantified cells, the resultant computer simulations produced highly different outcomes (Supplementary Fig. S10). We therefore decided to use a simplified approach in which we attempted to generally describe the band formation process and determine key properties that lead to robust separation. We thus created a “caricature” dataset, which we based on the average values obtained from our experiments (Supplementary Table 5). We then attempted to simulate the band formation by implementing two temporal phases: i) separation, where a large difference of the microtubule dynamic instability parameters between band and gap regions is applied (default 5x based on average *G*-ratio of 4.84 during fastest separation) and ii) maintenance, where a much smaller difference is applied (default 2x, based on average *G*-ratio of 2.26 during slowest separation). Although band-associated *G*-values fluctuated in some cells (Supplementary Fig. S5-S8 and S10), these fluctuations lacked a clear bias and therefore we kept these values constant at a level representative of “early”, i.e. prior to band separation, microtubule arrays in the simulations. From this baseline, we increased the catastrophe rate *r*_cat_ in the gaps by an integer factor. The resulting differences in *G*-values were similar to the large differences observed in our experimental data. As a key measure, we monitored the ‘*degree of separation*’ over time, which is the ratio of microtubule densities in bands versus gaps. However, we were only able to obtain a very low degree of separation using these parameters (Fig. 2b,e).

**Figure 2.**
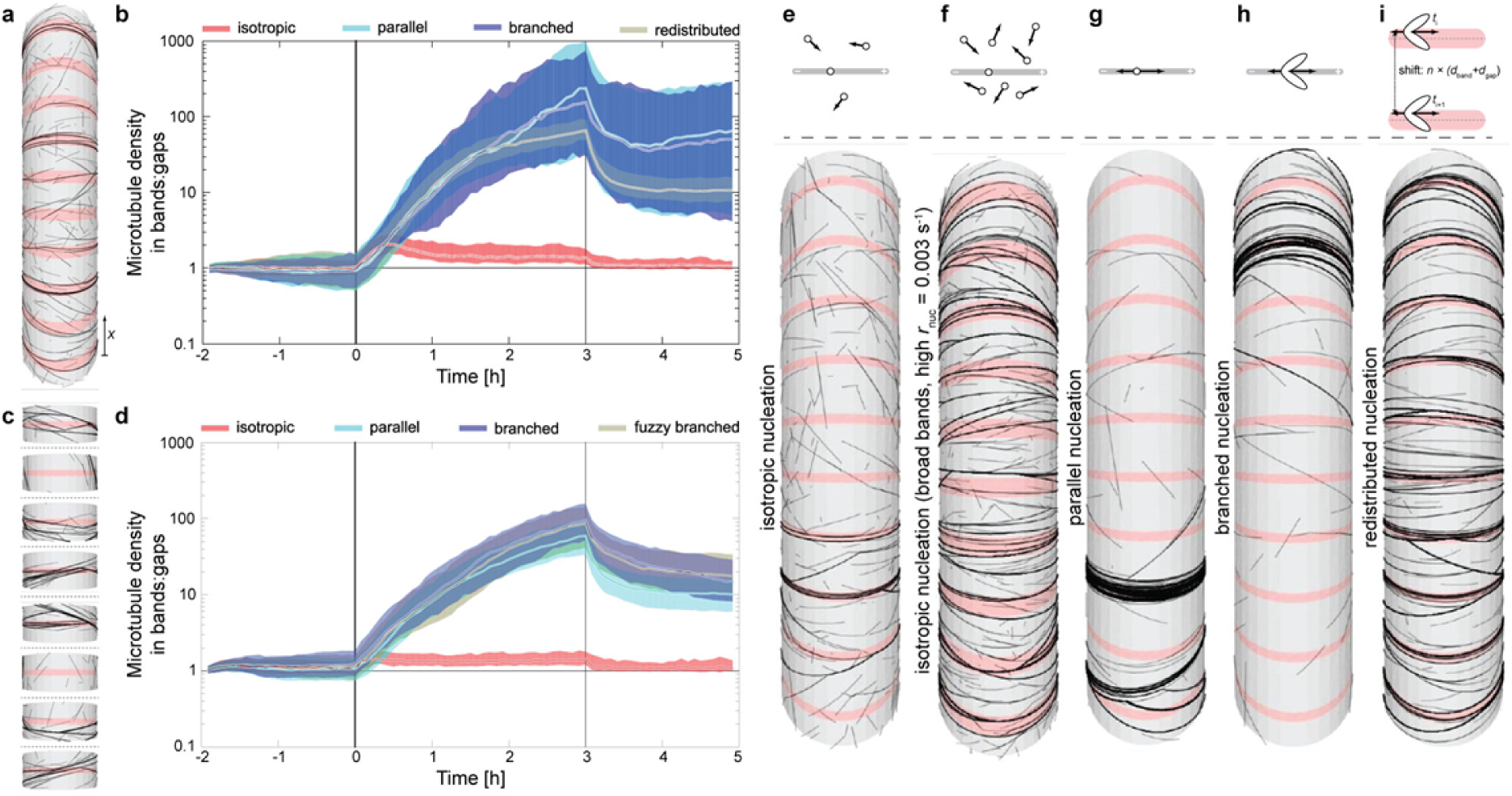
Simulation of dynamic microtubules reproduce patterning into bands and gaps. **a**, Design of the simulated cells: Dynamic microtubules (black lines) exist on a cylindrical surface with a diameter of 15 µm and a length of 60 µm. Ten 1 µm wide bands (pink) separated by 5 µm wide gaps were distributed over the cell surface. The separation phase runs from 0 to 180 minutes, followed by a 120 minutes maintenance phase. **b**, Ratio of microtubule density in bands and gaps of 10-banded cells for four different nucleation modes. Vertical lines divide initiation, separation and maintenance regimes. Median, and 16^th^-to-84^th^ percentile range from >100 independent simulations are shown for each nucleation mode. **c**, Alternative design of the simulated cells: single-banded cells including their gap environment with a length of 6 µm. All other parameters remained as in (A). **d**, Ratio of microtubule density in bands and gaps of single-banded cells for four different nucleation modes. Vertical lines indicate initiation, separation and maintenance regimes. Median, and 16^th^-to-84^th^ percentile range for >100 composite cells comprised of 10 randomly chosen single-band cells from a collection of >2000 independent single-banded simulations are shown for each nucleation mode. **e-i**, Top row of panels shows schematic representations of the five different nucleation modes: isotropic (e), isotropic with higher nucleation rate (f), parallel (g), branched (h) and redistributed nucleation (i). A microtubule is depicted as a grey rod with plus- and minus-ends. Arrows indicate the direction of a polymerizing microtubule from a nucleation site. Elliptic areas depict polymerization bias for branched nucleation of approx. 35 degrees relative to the microtubule axis. Redistributed nucleation involves register of the relative position of a nucleation site to the nearest band (pink bar) followed by a shift of a random integer times the length of the repeating unit (*d*_band_ + *d*_gap_) leading to global redistribution of the nucleation sites. Lower row of panels shows snapshots of representative simulations for each nucleation mode.

As indicated above, *G* integrates microtubule dynamic instability parameters and nucleation rates. In our initial simulations, we assumed isotropic nucleation (i.e., from random points in the array with random orientations). Alterations in the rate of nucleation, e.g. increased with a factor of three, did not markedly change the degree of separation (Fig. 2f). As microtubule nucleation occurs from existing microtubules in diffuse array configurations (Chan et al. 2009), we implemented such behavior in the simulations, i.e. microtubule nucleation becomes associated with microtubule bands during the separation phase. The corresponding simulations led to an increased degree of separation (Figs. 2b,g,h). Here, the relative nucleation angles – parallel (default; new microtubules are nucleated on existing ones in parallel orientations) or branched (similar to parallel with an additional option to branch off at 35 degree) – had little impact.

The strong effect of microtubule-bound nucleation on the degree of separation (Supplementary Information and Fig. S11) can be understood as a positive feedback loop: a local increase in microtubule density attracts more nucleation complexes, resulting in a further increase of the local microtubule density (Deinum et al. 2011). In our simulations, the microtubule density-based competition for nucleation complexes is global, i.e., over the whole cell length, which typically resulted in the over-amplification of a single or a few dominant bands (Fig. 2g,h and Supplementary Fig. S10). This unrealistic behavior indicated that *in planta*, the competition for nucleation complexes is local. To exclude global competition in our simulations, we continued with either short cells comprising only one band and then re-constituted the full-length cells from sets of ten single-banded cells (Fig. 2c, Supplementary Information), or we artificially re-distributed the nucleation sites randomly retaining their distance to the center of the nearest band (Fig. 2i). Both cases showed similar behavior corroborating the validity of the approach (Figs. 2b,d,i). These results suggest that band formation is restricted to a regime where the nucleation of microtubules preferentially occurs from the emerging bands and that nucleation complexes are locally recruited to the bands. In addition, the model allowed us to explore effects of modifying the *G* and *τ* parameters (see Supporting Information and Supplementary Fig. S12 and S13).

### Microtubule nucleation complexes are evenly distributed across microtubule bands

To investigate how our simulation predictions of nucleation distribution compared with microtubule nucleation in trans-differentiating cells, we crossed the GFP-labelled gamma-tubulin complex protein (GCP)3 (Nakamura et al. 2010) with the mCHERRY-TUA5 VND7 background. We found that GCP3 foci coincided with cortical microtubules in agreement with Nakamura et al. (2010), leading to an even distribution of GCP3 foci in non-induced cells and in the bands of induced cells (Figs. 3a-d). We measured the density of GCP3 foci and found that induced cells had significantly higher foci density in bands as compared to gaps (Fig. 3d), and compared to the even distribution observed at the cortex of non-induced cells (Figs. 3e,f). We did not find any statistical differences in the GCP3 density between neighboring bands (Fig. 3f). The GCP3 intensity gradient along the long axis of induced and non-induced cells was, furthermore, not significantly different from zero (Fig. 3g). These data are thus in good agreement with our simulation results.

**Figure 3.**
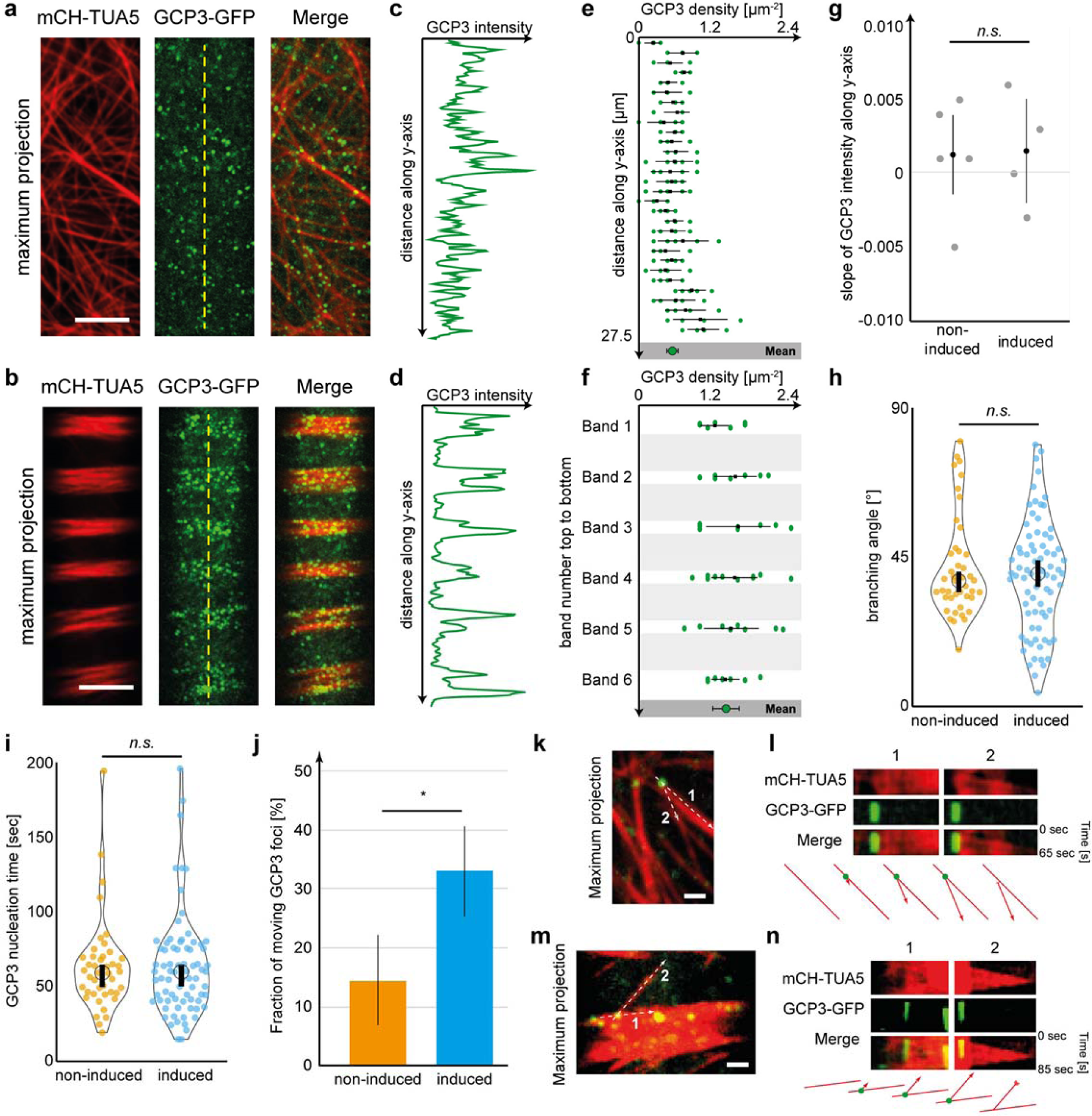
The microtubule-nucleating complex protein GCP3 distributes evenly across forming bands during xylem formation. **a-b**, Maximum projections of time series recordings of dual-labelled mCH-TUA5 GCP3-GFP in non-induced cells (a) and during mid stage of proto-xylem formation (b). Scale bar = 5 µm. **c-d**, Intensity scans along the dashed lines in (a) and (b) revealing a random cortical distribution of GCP3 foci in non-induced cells (c) and recruitment of GCP3 foci to forming microtubule bands in mid stages of proto-xylem formation (d). **e**, Cortical GCP3 density measured top to bottom along the *y*-axis of non-induced cells. Each green dot refers to a single density measurement, the black line and square represent mean ± s.d. for each section of the cortex. At the bottom (grey box), all measurements are pooled to an average GCP3 density (0.56 ± 0.19 foci per µm^2^, mean ± s.d., 155 measurements from 5 cells). **f**, GCP3 density measured top to bottom in microtubule bands in induced cells. Each dot refers to a single measurement; the black line and square represent mean ± s.d. for each band. At the bottom (grey box), all measurements are pooled to an average GCP3 density (1.41 ± 0.43 foci per µm^2^, mean ± s.d., 56 measurements from 4 cells, *p* < 0.0001, Welch’s unpaired *t*-test). **g**, Slope of the GCP3 intensity gradient along *y*-axis. Non-induced and xylem-forming cells showed no statistically significant correlation indicating an even distribution of GCP3 foci (mean ± 95% confidence intervals; *p* > 0.91, Welch’s unpaired *t*-test). **h-i**, Branching angle of new microtubules and nucleation time of nucleating GCP3 foci located parent microtubules are identical in non-induced (orange dots) and induced cells (blue dots), respectively (means ± 95% confidence intervals, p > 0.37 and p > 0.97, for 43 and 76 new microtubules and GCP3s from 5 and 4 cells, respectively). **j**, Proportion of moving GCP3s in non-induced (orange) and induced (blue) cells. For a GCP3 to be classified as moving, the foci had to move by more than 100 nm within 5 consecutive frames (25 seconds). Mean ± s.d., 43 and 76 GCP3s from 4 non-induced and 5 induced cells, *p* > 0.0142, Welch’s unpaired *t*-test. **k-n**, Maximum projections of time-lapse recordings showing GCP3-induced new microtubules being nucleated from parent microtubules in non-induced cells (k) and induced cells (m). Dashed lines indicate parent (1) and new (2) microtubule region used to generate kymographs (l, n). Scale bar = 1 µm. Panels l and n show kymographs of stationary GCP3s during nucleation of new microtubules for non-induced (l) and induced (n) cells. Below: schematic representation of the nucleations shown in the kymographs. red: microtubules, green: GCP3.

We next investigated whether the GCP3s promote microtubule nucleation during trans-differentiation. We focused on microtubules at the edge of bands, as nucleation events here were readily observable as compared to microtubules located in the center of bands, and compared those with microtubule nucleation in non-induced cells. We found that new microtubules were branched from parent microtubules at angles of approx. 40 degrees in both induced and non-induced cells (Fig. 3h). Similarly, microtubule nucleation typically occurred from GCP3 foci that had dwelled on microtubules for on average 60 seconds (Fig. 3i). In non-induced cells, the vast majority of nucleating GCP3s were stationary (86 ± 8 % of all foci moved less than 100 nm within 25 seconds, mean ± s.d., 5 cells, Figs. 3j,k,l) whereas this was the case in only 67 ± 8 % of the cases in induced cells (Figs. 3j,m,n). Interestingly, in induced cells the remaining GCP3s displayed linear movements along microtubules in the bands (Supplementary Movies 7-9). These movements resulted in a larger variability of angles of branching microtubules during their polymerization as compared to that in non-induced cells (Fig. 3h). Closer inspection revealed that the linear GCP3 movements were due to anti-parallel encounters of polymerizing, newly nucleated microtubules, followed by microtubule-microtubule contraction, i.e., microtubules sliding along each other, with velocities of 14 ± 8 nm/s (mean ± s.d., 35 GCP3s from 4 induced cells). This phenomenon was not observed in non-induced cells (Supplementary Movie 6). Together, these results indicate that our assumptions of microtubule nucleation in the simulations agree with how nucleation occurs during microtubule separation. Furthermore, anti-parallel microtubule-microtubule contractions seem to be an overlooked process that might assist in microtubule organization in bands.

### The alignment and separation speed of microtubule bands are impaired in the ***ktn1-2 mutant***

We next set out to investigate how defects in microtubule dynamics and organization influenced proto-xylem band separation. We therefore focused on the microtubule severing protein complex KATANIN (KTN), which is important for secondary wall production and microtubule alignment (Burk et al. 2001; Stoppin-Mellet et al. 2007). The KTN complex is a hexamer of KTN1-KTN80 heterodimers (Wang et al. 2017). The single copy of KTN1 is responsible for microtubule severing whereas four KTN80 isoforms confer targeting to microtubule crossovers. Consistent with a role of KTN in proto-xylem formation, KTN1 and at least one isoform of KTN80 were expressed and up-regulated approximately 8 hours after VND7 induction (Fig. 4a; Li et al. 2016b), which corresponds well with the time point when microtubules re-organize prior to band formation. To corroborate these data, we crossed a functional KTN1-GFP (Lindeboom et al. 2013), driven by its native promoter, into our mCHERRY-TUA5 marker line in the VND7 background. We found that in non-induced cells, KTN1-GFP co-localized dynamically with the microtubules (Fig. 4b). In induced cells, we found that KTN1-GFP strongly labelled the forming microtubule bands during the mid-stage of proto-xylem formation (Figs. 4c,d). These data confirm that KTN is associated with microtubules during proto-xylem formation and may be active during this process.

**Figure 4.**
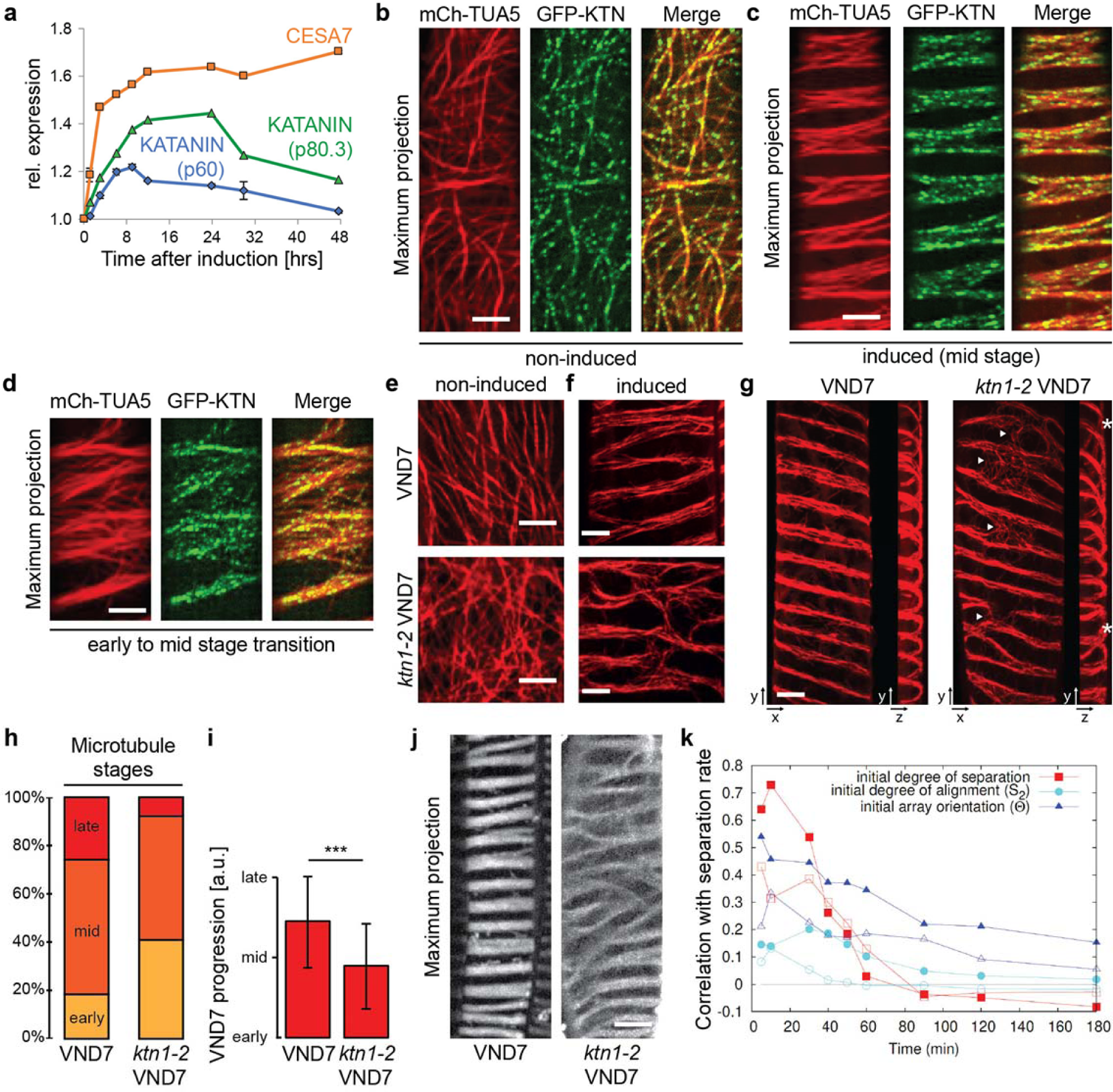
Microtubule Band Progression Depends on the Microtubule Organization Prior to Trans-differentiation. **a**, Expression of the KATANIN subunits p60 and p80.3, and a control gene for proto-xylem formation, secondary CESA7, is upregulated after VND7 induction. KATANIN subunits reach their maximum at 8 to 12 hrs post induction, which is the time when microtubule band formation occurs. Data normalized to the expression levels in non-induced control plants (data taken from Li et al., (2016b), mean ± sd, *n* = 3 replicates). **b-c**, Maximum projection of dual-labelled mCh-TUA5 and GFP-KTN seedlings in non-induced conditions (b) and mid stages of xylem formation (c). Scale bar = 10 µm. **d**, KTN-GFP localized majorly to microtubules in bands and the KTN-GFP signal was localized to bands during early and mid-stages of protoxylem formation. Scale bar = 10 µm. **e-f**, Time-average projections of mCherry-TUA5 microtubules in non-induced (e) and mid stages (f) of proto-xylem formation in wild-type (top) and *ktn1-2* mutant seedlings (bottom). Microtubule arrays in non-induced hypocotyl cells of wild-type seedlings showed better co-alignment compared to *ktn1-2* mutants. Induced seedlings showed compact microtubule bands and efficient separation in wild-type, whereas *ktn1-2* mutants showed microtubule bands that remained cross-linked. Scale bar = 5 µm. **g**, Maximum projections (left) and projections along the *x*-axis (right) of mCh-TUA5 VND7 in mid stages of xylem formation in wild-type and the *ktn1-2* background. In *ktn1-2*, microtubule bands remain connected for longer times (arrow heads) and the band spirals are more disordered (asterisks). Scale bar = 5 µm. **h**, Fraction of induced hypocotyl cells classified into early, mid, and late stages of xylem formation of mCh-TUA5 VND7 in wild-type and *ktn1-2* background. All investigated cells showed signs of trans-differentiation 18 hours after induction. The *ktn1-2* mutant is less progressed compared to wild-type. **i**, Relative progression of VND7 in wild-type and the *ktn1-2* background. Early-, mid-, and late-stage cells were given weights of 1, 2, and 3, respectively, allowing for the quantification of their average microtubule stage. Microtubule arrays in wild-type are significantly more progressed (2.1 ± 0.7, mean ± s.d., 174 cells from 9 seedlings) compared to the *ktn1-2* mutant (1.7 ± 0.6, 127 cells from 9 seedlings, *p* < 0.0001, Welch’s unpaired *t*-test). **j**, Maximum projection of the stained secondary walls of induced wild-type and *ktn1-2* seedlings 48 hours after induction. Scale bar = 10 µm. **k**, Correlations between increase of degree of separation (separation rate) since *T* = 0 and (i) initial degree of separation (ratio of band density to gap density; red), (ii) initial anisotropy (S2, cyan), and (iii) initial array orientation (*θ*, blue). All arrays are initiated with isotropic nucleation, followed by parallel nucleation during separation phase. Closed symbols: *P*_cat_ = 0.5 during entire simulation; open symbols: *P*_cat_ = 0.05 during initiation phase, and *P*_cat_ = 0.5 afterwards. *n* = 500 10-banded cells.

To obtain insight into how KTN affects proto-xylem microtubule band formation, we crossed a *ktn1-2* knockout mutant into our mCHERRY-TUA5 marker line in the VND7 background (Nakamura et al. 2010; Lindeboom et al. 2013). In agreement with previous work, we found that microtubules were more isotropically organized in the *ktn1-2* mutant (Fig. 4e and Supplementary Fig. S14). Nevertheless, *ktn1-2* mutants were clearly able to undergo microtubule separation upon VND7 induction (Figs. 4f,g). However, the resulting bands displayed defects in alignment and were less coherent than in the wild type. We observed that some microtubule bands did not separate well from each other and maintained bridging microtubule clusters (arrowheads in Fig. 4g). When counting the number of cells in early, mid, and late stages of proto-xylem formation (at 16 to 18 hours after induction) we found that the *ktn1-2* cells were substantially delayed (Figs. 4h,i). While the wild-type VND7 seedlings were on average between mid and late stages, the *ktn1-2* seedlings were not yet at mid stage (Fig. 4i). When staining the secondary cell walls of wild-type and *ktn1-2* seedling cells in the VND7 background we found similar misarrangements of cell wall bands in *ktn1-2* (Fig. 4j) that closely matched the aberrations observed for the microtubule array. These results indicate that KTN is not essential to drive microtubule band separation but does affect the fidelity and speed of this process.

### Array orientation and anisotropy impact separation speed *in silico*

To try to understand the KTN-induced changes in the band separation process, we included KTN-based microtubule severing at crossovers in our simulations (Deinum et al. 2017). We found that this katanin activity had little impact on the separation process, regardless of the type of nucleation mode (parallel, branched or “fuzzy branched”; Supplementary Fig. S15 and Supplementary Information). We next hypothesized that the main cause of delayed band formation in the *ktn1-2* mutant could be the impaired formation of well-aligned, anisotropic arrays prior and during the band separation. To investigate this, we altered the initiation phase to generate a wider range of initial array anisotropies and orientation angles at the start of the separation phase. To this end, we employed isotropic nucleation throughout the initiation phase. We also used two different values of *P_cat_*, which describes the induced catastrophe probability for large angle collisions, during the initiation phase. Here, we used *P_cat_* = 0.5 as a default, which still yields mostly well aligned arrays, and 10-fold reduced *P_cat_* (0.05), which predominantly yields poorly aligned arrays (Supplementary Fig. S16). We then computed the correlation coefficients between the separation rate (relative increase of the degree of separation) and (i) the degree of separation at T = 0 (the start of separation phase), (ii) the degree of alignment (*S_2_*; a measure of anisotropy; Deinum et al. 2017) at *T* = 0, and (iii) the array orientation (*Θ*) at multiple time points (Fig. 4k; scatter plots at *T* = 10 minutes in Supplementary Fig S16). We found that for both data sets, initial array orientation (blue triangles) correlated stronger with separation rate than anisotropy (cyan circles). The effect of orientation was, moreover, stronger for well-aligned arrays (blue vs. empty triangles). These findings suggest that fast separation requires both a transverse and anisotropic array. During the first 40-50 minutes, there was a strong correlation of separation rate with the degree of separation at the onset of the separation phase (red squares). This observation suggests that spontaneous density fluctuations of the microtubule array at *T* = 0 are quickly enhanced and thus drive the reorientation of the overall array into the transverse state. These results provide explanations why the *ktn1-2* mutant – which shows lower anisotropy, and more disperse microtubule orientations across the cortex – displayed slower microtubule separation rates than the wild-type array. The observation that the microtubule bands in the *ktn1-2* mutant remained connected for longer periods also suggests that the KTN function plays a role in defining band-gap boundaries.

## DISCUSSION

Microtubule patterning templates the deposition of cell wall components, typified during xylem differentiation. Patterned secondary wall deposition is best understood during pit formation; a specific patterned cell wall type of meta-xylem with small pit regions lacking secondary walls. Using VND6, a master regulator of meta-xylem development, future pit regions were found to be marked by the active form of ROP11, which recruits the microtubule-associated protein (MAP) MIDD1 to promote microtubule disassembly, to the plasma membrane (Oda et al. 2010; Oda and Fukuda 2012a). MIDD1 in turn recruits KIN13A, a microtubule motor protein that depolymerizes microtubules (Oda and Fukuda 2013). A Turing-like reaction-diffusion mechanism (Turing 1952) is suggested to initiate patterning of ROP11 in meta-xylem formation, in which ROPGEF4 and ROPGAP3 may act as activator and inhibitor, respectively (Oda and Fukuda 2012b). Alternatively, the difference in diffusion constants between active and inactive ROP11 may drive the Turing mechanism (Jacobs et al. 2019). Both mechanisms would explain how ROP11-induced microtubule-free regions are established to spatially control cell wall deposition (Oda and Fukuda 2012a; Oda and Fukuda 2013). These studies provided snapshots of microtubule pattern formation during meta-xylem differentiation, but the underlying microtubule array behaviour has remained unclear. ROP11-mediated patterning does not impact proto-xylem formation, since neither the presence of a constitutively active or dominant negative version of ROP11, nor the absence of single ROPGEFs and ROPGAPs, alter the spacing between secondary cell wall thickenings in the proto-xylem (Oda and Fukuda 2012a; Nagashima et al. 2018). The molecular contributors to proto-xylem patterning are therefore not defined. Nonetheless, ROP1 and ROP7 are putative candidates since these GTPases are upregulated after VND7 induction (Yamaguchi et al. 2011).

Our study reveals that microtubule behaviour changes spatiotemporally during proto-xylem trans-differentiation. There are clear similarities in proto- and meta-xylem formation as cortical regions with a reduced microtubule density show absence of wall thickenings. In addition, we observed unchanged *v*_+_ and *v*_-_, but a temporal increase in *r*_cat_, in microtubule gaps, similar to that in microtubules of developing pits (Oda et al. 2010). This similarity might indicate that microtubule disassembly in gaps and pits is driven by analogous mechanisms, perhaps via MIDD1 and KIN13A as both are upregulated upon VND7 induction (Yamaguchi et al. 2011). Contrary to the reduced microtubule rescue rates observed at pits (Oda et al. 2010), we found an increase in the microtubule rescue rates in the band gaps. These results indicate that some common but perhaps also diverging functions drive proto- and meta-xylem pattern formation.

We observed that bands formed simultaneously across the trans-differentiating cells, indicating that inter-band spacing and orientation is dictated by a cell-wide pattern that has formed before, or during the onset of microtubule rearrangements. This process, in conjunction with the ability to merge incoherent bands, occurred rapidly and appears crucial for accurate cell wall patterning. A modelling study on ROP patterning suggested that microtubule arrays, which form a barrier to ROP diffusion, cannot change the orientation of an existing pattern. This indicates that microtubules need to be oriented transversely prior to the onset of band separation (Jacobs et al. 2019).

Our simulation approach using a compiled parameter set revealed that spatiotemporal differential microtubule dynamics (*G*-values) and lifetimes (*τ*) are two key factors defining the degree and speed of band separation (see Supporting Information). When comparing this finding with the experimental time series, we found that increases in the separation rate indeed occurred at time points when the ratios of *G*_gap_/*G*_band_ and *τ*_band_/ *τ*_gap_ peaked. In some cases, such peaks were observed shortly before an increase in separation rate, indicating that microtubule breakdown in the gaps is a limiting step in band separation. Furthermore, our simulations showed that band separation occurred efficiently not only when nucleation rates were reduced in the gaps, but on the contrary, the nucleation rates had to be increased in the bands and a mechanism had to be in place that evenly distributed active GCP3 complexes over the microtubule bands. This indicates that microtubule bands are also actively supported during proto-xylem formation.

In meta-xylem, microtubules confine the gap regions by acting as diffusion barriers for active ROP11 (Oda and Fukuda 2012a; Sugiyama et al. 2017). Proteins that influence the properties of microtubule arrays might thus play an important role in fine-tuning the shape and fidelity of gaps in proto-xylem. To exemplify this mechanism, we focused on KATANIN and showed that it exerts a function in proto-xylem formation. While *ktn1-2* mutants produced microtubule bands, their shape, relative positioning and timely appearance were altered. In our simulations, we started with arrays that had a similar orientation as the orientation of the band region. This explains why the severing rate had little influence on the separation of well-ordered transverse arrays *in silico* as those arrays are in a “perfect” starting configuration with hardly any cross-overs for KATANIN to sever. When we instead investigated less-ordered arrays, we found that array orientation, anisotropy, and spontaneous microtubule density fluctuations were important determinants of separation rate. These parameters are altered in the *ktn1-2* mutant and we propose that KATANIN may support anisotropic, transverse microtubule patches prior to band formation. These findings further support the idea that gap regions are actively maintained through microtubule-mediated barriers.

KATANIN has also been implicated in releasing microtubules from nucleating GCPs (Nakamura et al. 2010). This may provide an explanation for the band-to-band microtubule cross-overs in the *ktn1-2* mutant. The absence of KATANIN is likely to lead to microtubules growing into existing gap regions at angles of approx. 40 degrees, which despite the increased microtubule catastrophe rates in the gaps, may cause reduced definition of the band-gap boundary. Perhaps such reduction could stabilize some aberrant microtubules to form the microtubule band-to-band cross-overs. The maintenance of microtubule bands is also driven by the ability to merge bands that are incoherent to the periodic band patterns. MAP70-5 may be a factor involved in delimiting microtubule bands, since it localises to the periphery of microtubule bands (Pesquet et al. 2010), perhaps with the aid of other MAPs, such as MAP65-8 and MAP70-1, that are expressed during treachery element formation (Derbyshire et al. 2015).

In summary, our study provides substantial new insight into how microtubules are re-arranged to sustain cell wall patterning. The simulation approaches may be extended to further predict microtubule-related functions to spatio-temporally control microtubule band separation, and could also allow for similar assessments during other types of wall patterning.

## ACKNOWLEDGEMENT

Kris van‘t Klooster was supported by the SRON space research program with project number GO-MG/15, which is financed by the Netherlands Organization for Scientific Research (NWO). We would like to thank the Wageningen Light Microscopy Centre (WU) and the Biological Optical Microscopy Platform (BOMP, University of Melbourne) for the use of their facilities and Bela Mulder for helpful discussions. Staffan Persson was supported by R@MAP Professor Funds at University of Melbourne and an ARC DP and FT grants (DP190101941; FT160100218).

## SUPPLEMENTARY TABLES

**Supplementary Table 1.** Microtubule dynamic instability parameters for cell 1

| Region | Time [min] | $v_+$ [$\mu\text{m}/\text{min}$] ± s.d. | $v_-$ [$\mu\text{m}/\text{min}$] ± s.d. | Number microtubule ends | $r_{\text{cat}}$ [/MT/s] | $r_{\text{res}}$ [/MT/s] | $G_{\text{gap}}/G_{\text{band}}$ |
| --- | --- | --- | --- | --- | --- | --- | --- |
| Band Gap | 0 | 0.064±0.019 | -0.123±0.050 | 282 | 0.002 | 0.001 | 1.046 |
| | | 0.065±0.019 | -0.123±0.052 | 374 | 0.002 | 0.001 | |
| Band Gap | 30 | 0.069±0.024 | -0.119±0.038 | 188 | 0.004 | 0.002 | 1.034 |
| | | 0.073±0.029 | -0.112±0.037 | 213 | 0.004 | 0.003 | |
| Band Gap | 60 | 0.070±0.023 | -0.102±0.038 | 268 | 0.003 | 0.001 | 0.896 |
| | | 0.070±0.025 | -0.107±0.041 | 347 | 0.002 | 0.001 | |
| Band Gap | 90 | 0.072±0.023 | -0.102±0.045 | 176 | 0.004 | 0.003 | 1.654 |
| | | 0.068±0.023 | -0.105±0.039 | 119 | 0.006 | 0.004 | |
| Band Gap | 120 | 0.072±0.020 | -0.099±0.037 | 255 | 0.003 | 0.002 | 3.272 |
| | | 0.073±0.019 | -0.110±0.069 | 72 | 0.009 | 0.006 | |
| Band Gap | 150 | 0.075±0.025 | -0.107±0.039 | 342 | 0.002 | 0.001 | 4.283 |
| | | 0.078±0.025 | -0.116±0.046 | 71 | 0.009 | 0.004 | |
| Band Gap | 180 | 0.073±0.021 | -0.090±0.034 | 173 | 0.004 | 0.003 | 7.750 |
| | | 0.083±0.026 | -0.113±0.043 | 27 | 0.027 | 0.012 | |
| Band Gap | 210 | 0.080±0.027 | -0.089±0.033 | 183 | 0.004 | 0.003 | 4.460 |
| | | 0.080±0.027 | -0.105±0.046 | 43 | 0.013 | 0.007 | |

**Supplementary Table 2.** Microtubule dynamic instability parameters for cell 2

| Region | Time [min] | $v_+$ [$\mu\text{m}/\text{min}$] ± s.d. | $v_-$ [$\mu\text{m}/\text{min}$] ± s.d. | Number microtubule ends | $r_{\text{cat}}$ [/MT/s] | $r_{\text{res}}$ [/MT/s] | $G_{\text{gap}}/G_{\text{band}}$ |
| --- | --- | --- | --- | --- | --- | --- | --- |
| Band Gap | 0 | 0.038±0.013 | -0.073±0.025 | 288 | 0.0015 | 0.0008 | 0.995 |
| | | 0.038±0.013 | -0.072±0.027 | 403 | 0.0015 | 0.0008 | |
| Band Gap | 30 | 0.048±0.014 | -0.074±0.031 | 306 | 0.0010 | 0.0006 | 3.375 |
| | | 0.052±0.014 | -0.081±0.028 | 212 | 0.0034 | 0.0016 | |
| Band Gap | 60 | 0.050±0.013<br>0.053±0.019 | -0.073±0.031<br>-0.079±0.030 | 372<br>215 | 0.0006<br>0.0024 | 0.0003<br>0.0009 | 3.950 |
| Band Gap | 90 | 0.043±0.011<br>0.047±0.011 | -0.070±0.025<br>-0.075±0.030 | 165<br>96 | 0.0008<br>0.0022 | 0.0003<br>0.0006 | 2.477 |
| Band Gap | 120 | 0.041±0.012<br>0.042±0.011 | -0.079±0.027<br>-0.090±0.031 | 196<br>37 | 0.0008<br>0.0054 | 0.0003<br>0.0032 | 5.746 |
| Band Gap | 150 | 0.044±0.010<br>0.043±0.009 | -0.088±0.022<br>-0.101±0.051 | 145<br>23 | 0.0019<br>0.0058 | 0.0011<br>0.0029 | 3.247 |
| Band Gap | 180 | 0.028±0.005<br>0.027±0.007 | -0.074±0.027<br>-0.084±0.030 | 115<br>31 | 0.0021<br>0.0046 | 0.0013<br>0.0022 | 2.380 |

**Supplementary Table 3.** Microtubule dynamic instability parameters for cell 3

| Region | Time [min] | $v_+$ [$\mu\text{m}/\text{min}$]<br>± s.d. | $v_-$ [$\mu\text{m}/\text{min}$]<br>± s.d. | Number microtubule ends | $r_{\text{cat}}$ [MT/s] | $r_{\text{res}}$ [MT/s] | $G_{\text{gap}}/G_{\text{band}}$ |
| --- | --- | --- | --- | --- | --- | --- | --- |
| Band Gap | 0 | 0.031±0.010<br>0.030±0.012 | -0.056±0.027<br>-0.059±0.025 | 180<br>268 | 0.0007<br>0.0011 | 0.0002<br>0.0007 | 1.651 |
| Band Gap | 30 | 0.035±0.014<br>0.034±0.012 | -0.055±0.026<br>-0.053±0.022 | 164<br>75 | 0.0007<br>0.0040 | 0.0003<br>0.0021 | 6.045 |
| Band Gap | 60 | 0.034±0.008<br>0.037±0.009 | -0.042±0.016<br>-0.048±0.015 | 157<br>46 | 0.0005<br>0.0038 | 0.0003<br>0.0012 | 7.455 |
| Band Gap | 90 | 0.037±0.013<br>0.036±0.010 | -0.057±0.027<br>-0.065±0.026 | 164<br>71 | 0.0003<br>0.0020 | 0.0002<br>0.0005 | 7.212 |
| Band Gap | 120 | 0.038±0.013<br>0.045±0.018 | -0.057±0.025<br>-0.063±0.024 | 96<br>52 | 0.0007<br>0.0032 | 0.0003<br>0.0015 | 3.904 |
| Band Gap | 150 | 0.033±0.011<br>0.037±0.013 | -0.058±0.026<br>-0.060±0.020 | 91<br>103 | 0.0006<br>0.0015 | 0.0002<br>0.0006 | 2.068 |
| Band Gap | 180 | 0.039±0.010<br>0.039±0.014 | -0.050±0.020<br>-0.047±0.019 | 96<br>83 | 0.0007<br>0.0018 | 0.0003<br>0.0003 | 3.127 |
| Band Gap | 210 | 0.043±0.015<br>0.043±0.015 | -0.058±0.029<br>-0.064±0.025 | 73<br>83 | 0.0012<br>0.0028 | 0.0004<br>0.0009 | 2.356 |

**Supplementary Table 4.** Microtubule dynamic instability parameters for cell 4

| Region | Time [min] | $v_+$ [$\mu\text{m}/\text{min}$]<br>± s.d. | $v_-$ [$\mu\text{m}/\text{min}$]<br>± s.d. | Number microtubule ends | $r_{\text{cat}}$ [MT/s] | $r_{\text{res}}$ [MT/s] | $G_{\text{gap}}/G_{\text{band}}$ |
| --- | --- | --- | --- | --- | --- | --- | --- |
| Band Gap | 0 | 0.058±0.020<br>0.056±0.019 | -0.080±0.027<br>-0.100±0.047 | 75<br>178 | 0.0027 | 0.0016 | 1.041 |
| Band Gap | 30 | 0.055±0.020<br>0.065±0.033 | -0.095±0.041<br>-0.095±0.043 | 193<br>167 | 0.0010 | 0.0004 | 1.331 |
| Band Gap | 60 | 0.055±0.018<br>0.058±0.026 | -0.081±0.032<br>-0.091±0.039 | 203<br>74 | 0.0013 | 0.0006 | 2.093 |
| Band Gap | 90 | 0.054±0.020<br>0.052±0.017 | -0.086±0.030<br>-0.083±0.037 | 210<br>84 | 0.0015 | 0.0009 | 3.021 |
| Band Gap | 120 | 0.057±0.019<br>0.055±0.017 | -0.084±0.034<br>-0.083±0.039 | 277<br>124 | 0.0008 | 0.0005 | 4.893 |
| Band Gap | 150 | 0.057±0.019<br>0.053±0.018 | -0.074±0.030<br>-0.080±0.037 | 204<br>182 | 0.0012 | 0.0005 | 2.001 |
| Band Gap | 180 | 0.059±0.021<br>0.060±0.025 | -0.098±0.046<br>-0.096±0.046 | 162<br>136 | 0.0014 | 0.0006 | 1.109 |
| Band Gap | 210 | 0.058±0.016<br>0.059±0.020 | - 0.083±0.038<br>-0.086±0.040 | 172<br>129 | 0.0013 | 0.0004 | 1.652 |
| Band Gap | 240 | 0.057±0.017<br>0.064±0.029 | -0.071±0.030<br>-0.073±0.028 | 153<br>77 | 0.0011 | 0.0006 | 2.958 |
| Band Gap | 270 | 0.062±0.035<br>0.060±0.020 | -0.078±0.038<br>-0.097±0.038 | 180<br>80 | 0.0009 | 0.0003 | 3.537 |
| Band Gap | 300 | 0.069±0.030<br>0.060±0.022 | -0.085±0.038<br>-0.100±0.052 | 165<br>91 | 0.0008 | 0.0004 | 3.858 |
| Band Gap | 330 | 0.066±0.023<br>0.066±0.027 | -0.089±0.040<br>-0.105±0.027 | 124<br>63 | 0.0015 | 0.0006 | 2.438 |

**Supplementary Table 5.** Microtubule dynamic instability parameters for caricature dataset

| Region | Phase | Time [min] | $v_+$<br>[$\mu\text{m}/\text{min}$]<br>± s.d. | $v_-$<br>[$\mu\text{m}/\text{min}$]<br>± s.d. | $r_{\text{cat}}$<br>[MT/s] | $r_{\text{res}}$<br>[MT/s] | $G_{\text{gap}}/$<br>$G_{\text{band}}$ |
| --- | --- | --- | --- | --- | --- | --- | --- |
| Band Gap | Initiation | -120 | 0.050 | -0.080 | 0.0016 | 0.001 | 1 |
| Band Gap | Separation | 0 | 0.050 | -0.080 | 0.0016 | 0.001 | 4.84 |
| Band Gap | Mintenance | 180-300 | 0.050 | -0.080 | 0.0016 | 0.001 | 2.26 |

## SUPPLEMENTARY FIGURES

**Supplementary Fig. S1.**
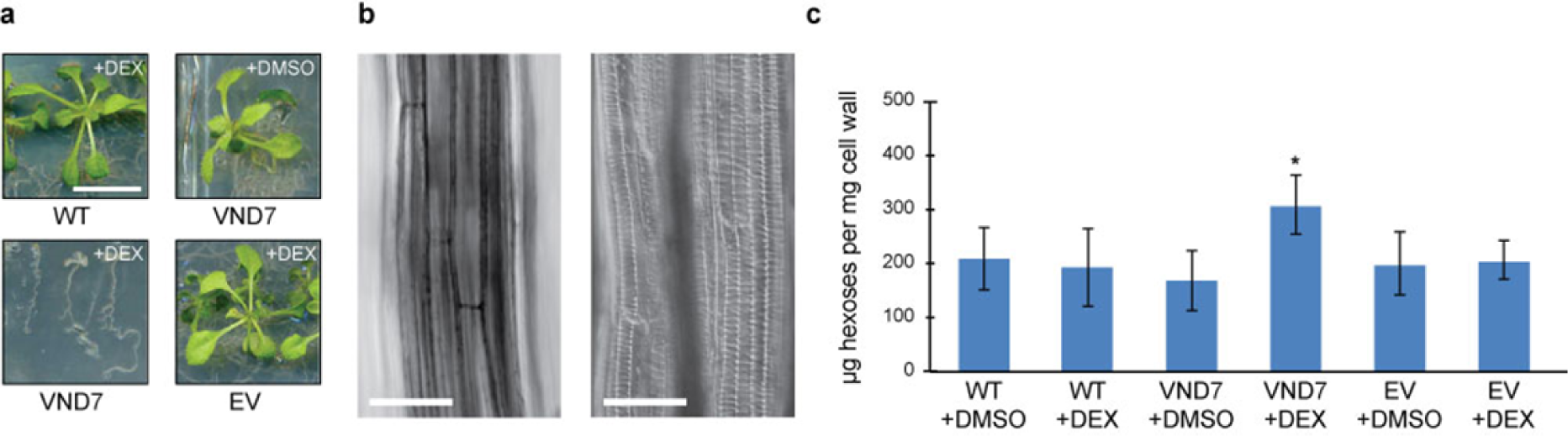
The VND7-inducible system drives secondary wall deposition in hypocotyl cells. **a**, Images of 10-day old light-grown wild-type, VND7 and empty-vector seedlings treated with 10 µM DEX and DMSO, respectively, for 72 hours. Scale bar = 1 cm. **b**, Differential interference contrast images of 10-day old dark-grown hypocotyls of wild-type (left) and VND7 (right). Scale bar = 100 µm. **c**, VND7-induced seedlings contain significantly more hexose sugars in their cell walls (309 ± 55 µg per mg cell wall material) compared to wild-type (208 ± 58 µg) and empty-vector seedlings (199 ± 59 µg; mean ± sd, *N* = 3 biological replicates with 3 technical replicates, \**p* ≤ 0.05, Welch’s unpaired *t*-test).

**Supplementary Fig. S2.**
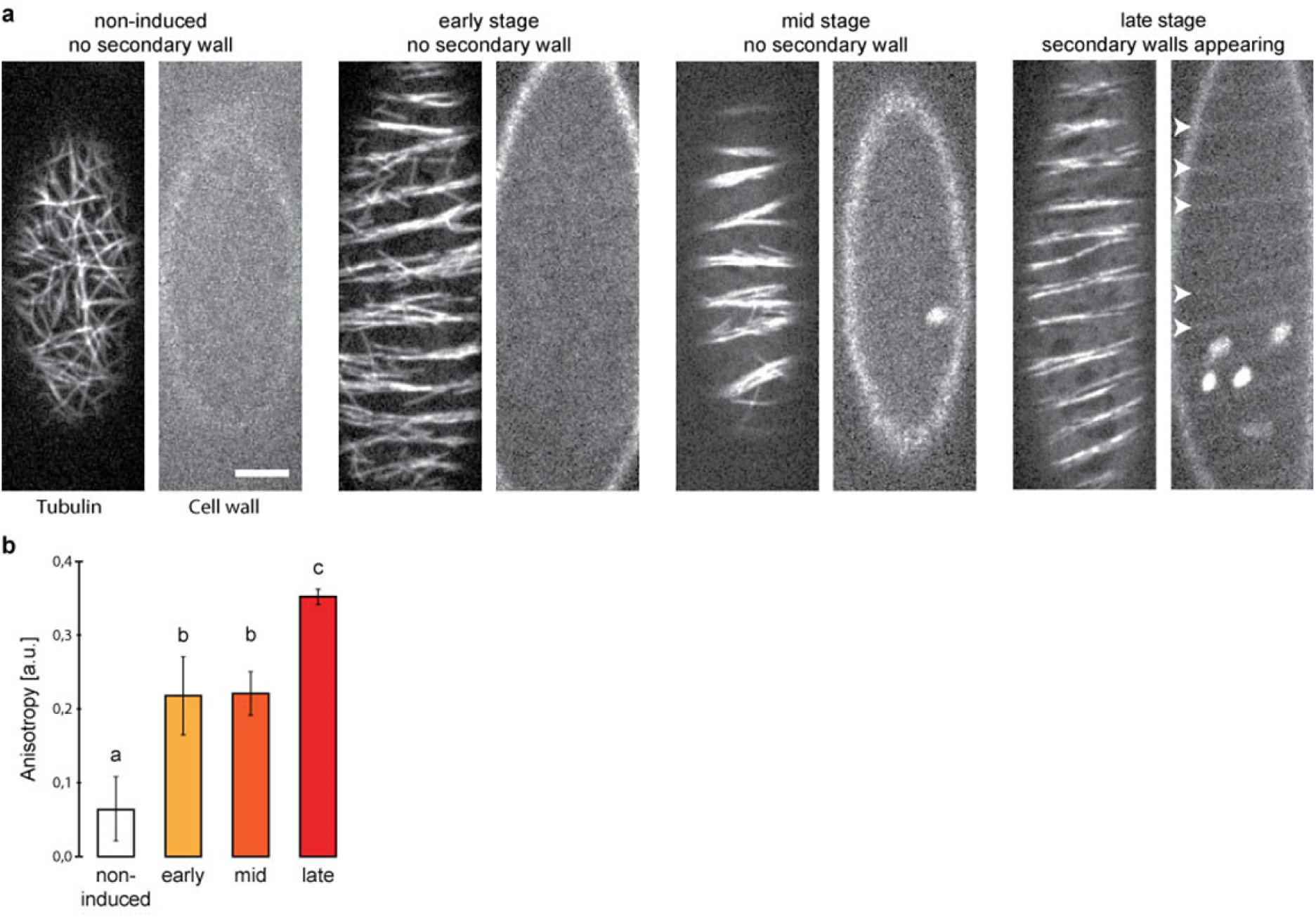
The VND7-inducible system drives cortical microtubule re-orientations. **a**, Representative stages of the VND7-driven secondary wall program: microtubule rearrangements precede the deposition of secondary walls (arrows heads in late stage images, stained by Direct Red 23, Material & Methods). Scale bar = 5 µm. **b**, The anisotropy of the microtubule array increased 5-fold from an initial 0.06 ± 0.04 in non-induced cells to a final 0.35 ± 0.01 (mean ± s.d., 7 cells from 5 seedlings) for cells in the late stage of proto-xylem formation. Groups assigned by statistical differences based on Welch’s unpaired *t*-test (p < 0.05 between groups).

**Supplementary Fig. S3.**
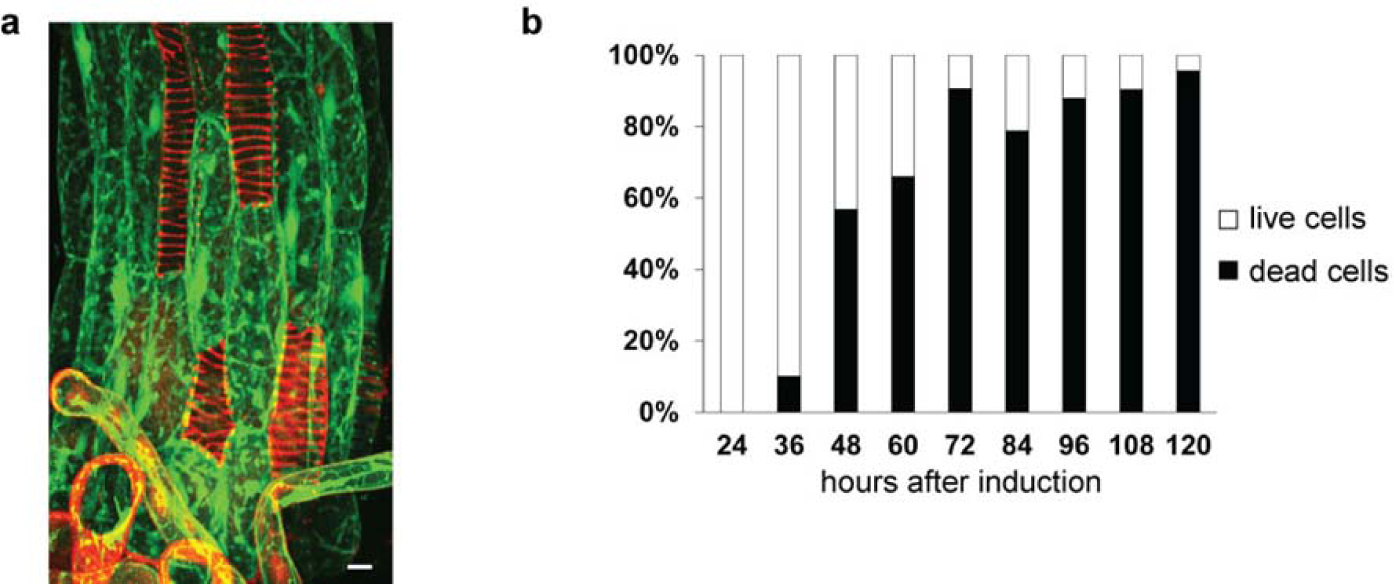
The VND7-inducible system leads to complete cell death after several days. **a**, VND7 induction leads to controlled cell death starting after 24 hours after induction. Cell death is assessed via the life-stain fluorescein diacetate (FDA), which is converted by intracellular esterase activity in living cells resulting in fluorescein fluorescence (green). No esterase activity is present in dead cells resulting in no staining. Propidium iodide (PI, red) stains the secondary cell wall thickenings. Scale bar = 10 µm. **b**, Time course of the fraction of cells undergoing programmed cell death after VND7 induction. Cells without FDA but with PI stain in (a) have completed the VND7 program and were scored as ‘dead’ cells (black). The majority of cells had died 120 hours after induction.

**Supplementary Fig. S4.**
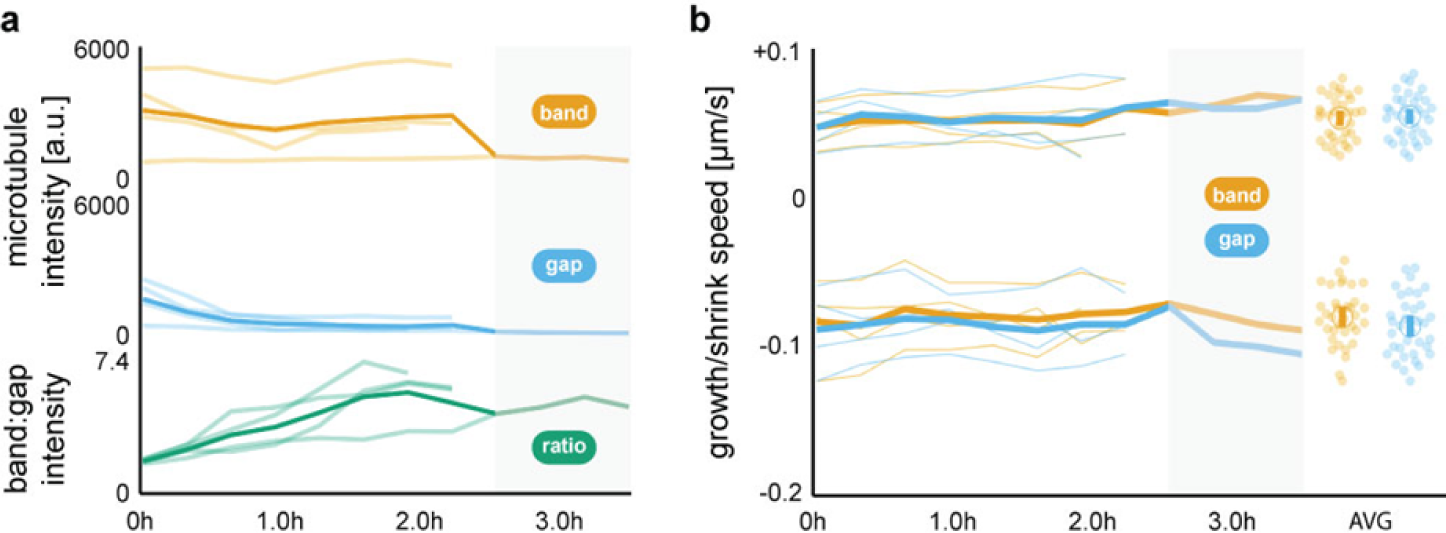
Quantification of microtubule intensity and dynamics in four independent time series’. **a**, Total microtubule intensity of four individual cells (pale lines) divided into band (orange) and gap regions (blue), and the microtubule intensity ratio between bands and gaps (green). The thickened line represent the time average of all measurements. **b**, Growth speed *v*_+_ and shrink speed *v*_-_ of four individual cells (fine lines) divided into band (orange) and gap regions (blue). The thickened line represent the time average of all measurements. Scatter plots: Data pooled from all 35 time points of all four cells; means ± 95 % confidence intervals. No differences between growth and shrinkage speeds were found between bands and gaps. The shaded area in panel a and b indicates a period covered by only one dataset (Cell 4). Statistics: Welch’s unpaired *t*-test, ** p < 0.005.

**Supplementary Fig. S5.**
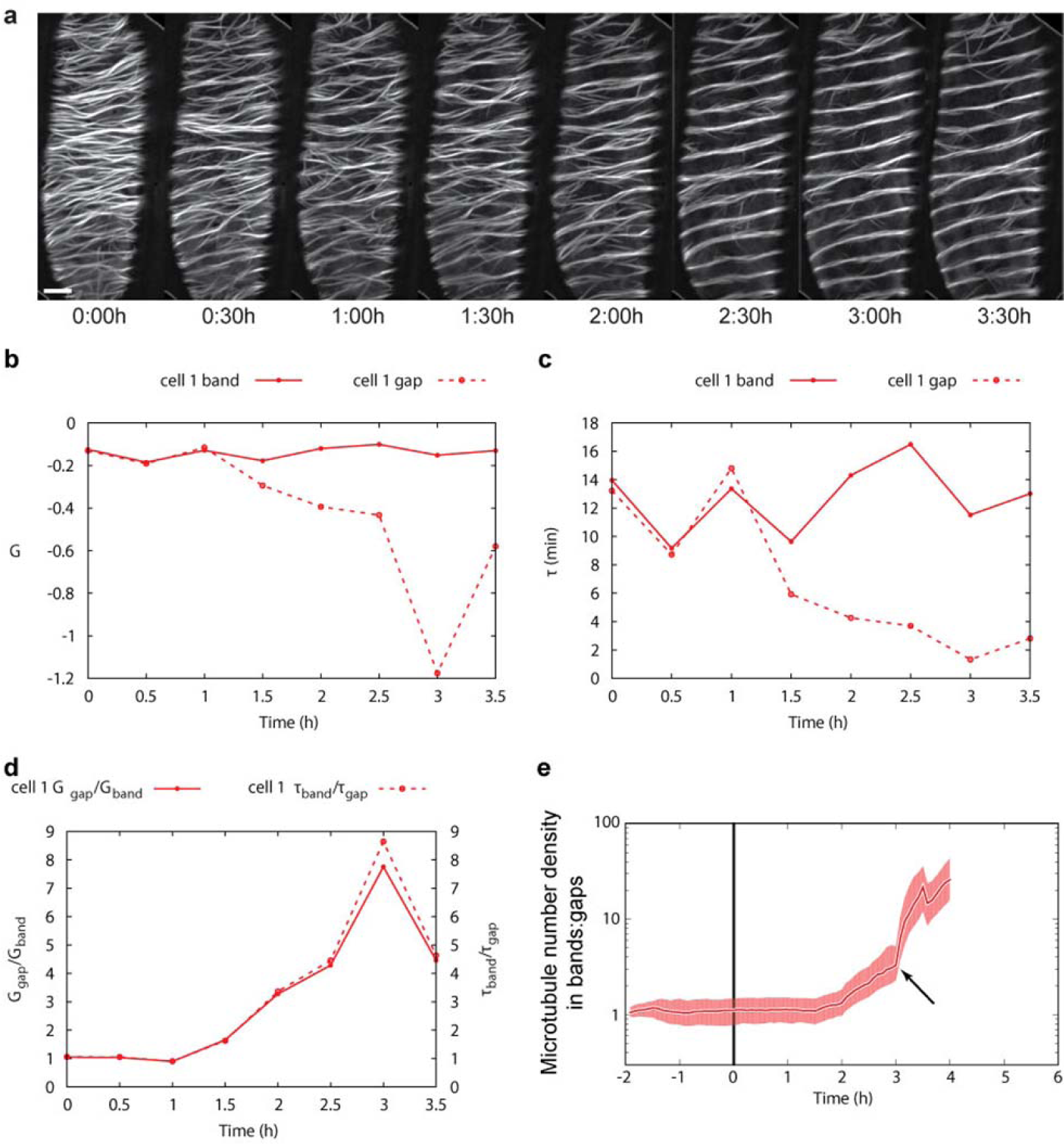
Microtubule re-orientations during the VND7-driven secondary wall program of cell 1. **a**, Average projections of YFP-labelled microtubules in an induced hypocotyl cell. Scale bar = 5 µm. **b-c**, Temporal development of the control parameter *G* (b) and the microtubule lifetime *τ* (c) in bands (solid line) and gaps (dashed line). Microtubules in the bands show more interactions and longer lifetimes than in the gaps. **d**, The temporal development of the ratio of *G*_gap_/*G*_band_ (solid line) and *τ*_band_*/ τ*_gap_ (dashed line) indicates that the number of microtubule-microtubule interactions and the microtubule lifetime needs to be sufficiently different between bands and gaps for separation to occur. **e**, Degree of separation measured as microtubule number density ratio between bands and gaps. Note that when the ratio of *G*_gap_/*G*_band_ and *τ*_band_*/ τ*_gap_ peaks at 3 h in (d) a strong increase in the separation rate is observed (arrow). Median ± 16 % and 84 % percentiles. *n* ≥ 100.

**Supplementary Fig. S6.**
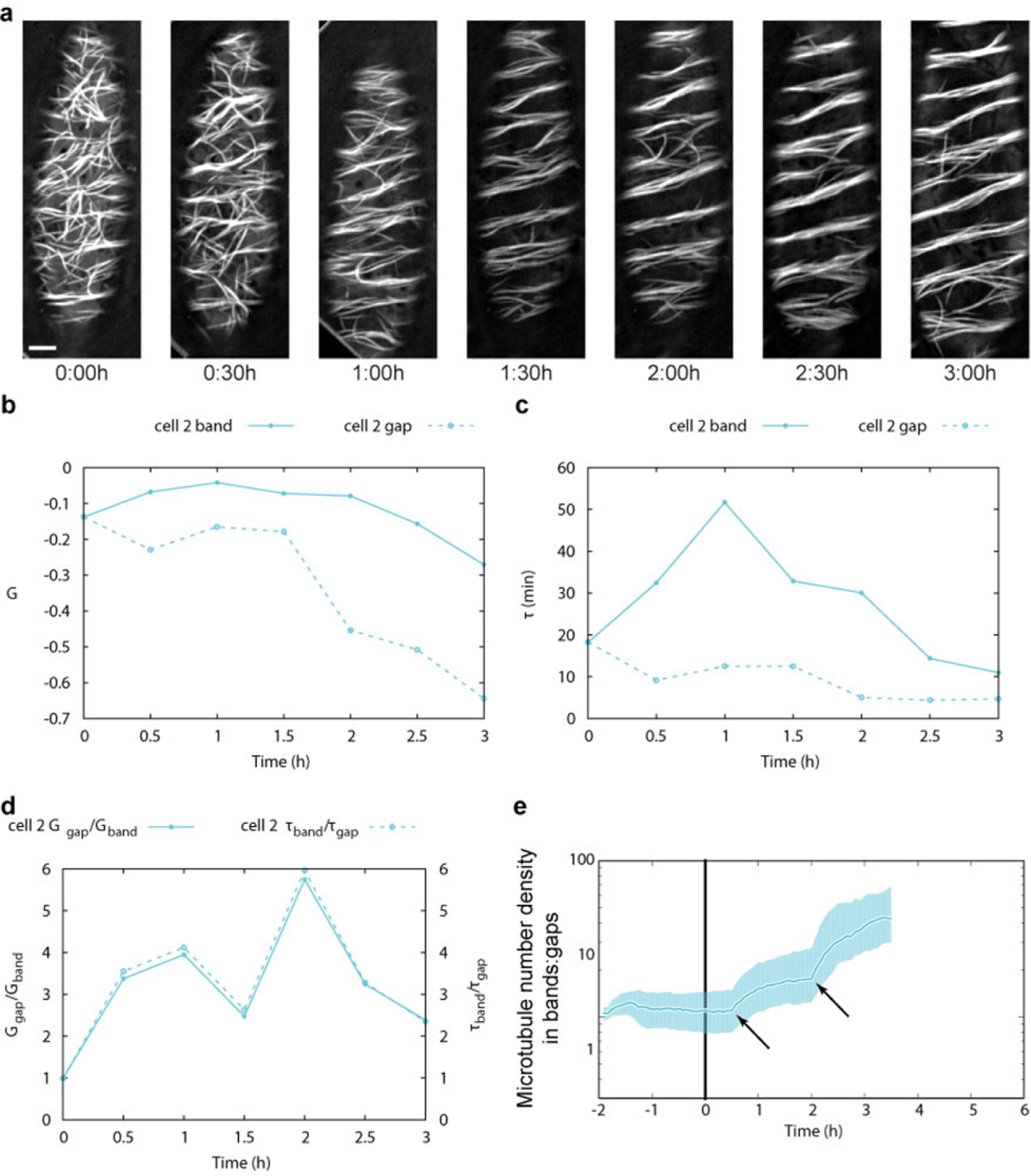
Microtubule re-orientations during the VND7-driven secondary wall program of cell 2. **a**, Average projections of YFP-labelled microtubules in an induced hypocotyl cell. Scale bar = 5 µm. **b-c**, Temporal development of the control parameter *G* (b) and the microtubule lifetime *τ* (c) in bands (solid line) and gaps (dashed line). Microtubules in the bands show more interactions and longer lifetimes than in the gaps. **d**, The temporal development of the ratio of *G*_gap_/*G*_band_ (solid line) and *τ*_band_*/ τ*_gap_ (dashed line) indicates that the number of microtubule-microtubule interactions and the microtubule lifetime needs to be sufficiently different between bands and gaps for separation to occur. **e**, Degree of separation measured as microtubule number density ratio between bands and gaps. Note that when the ratio of *G*_gap_/*G*_band_ and *τ*_band_*/ τ*_gap_ peaks at 0.5 h to 1 h and 2 h in (d) strong increases in the separation rate are observed (arrows). Median ± 16 % and 84 % percentiles. *n* ≥ 100.

**Supplementary Fig. S7.**
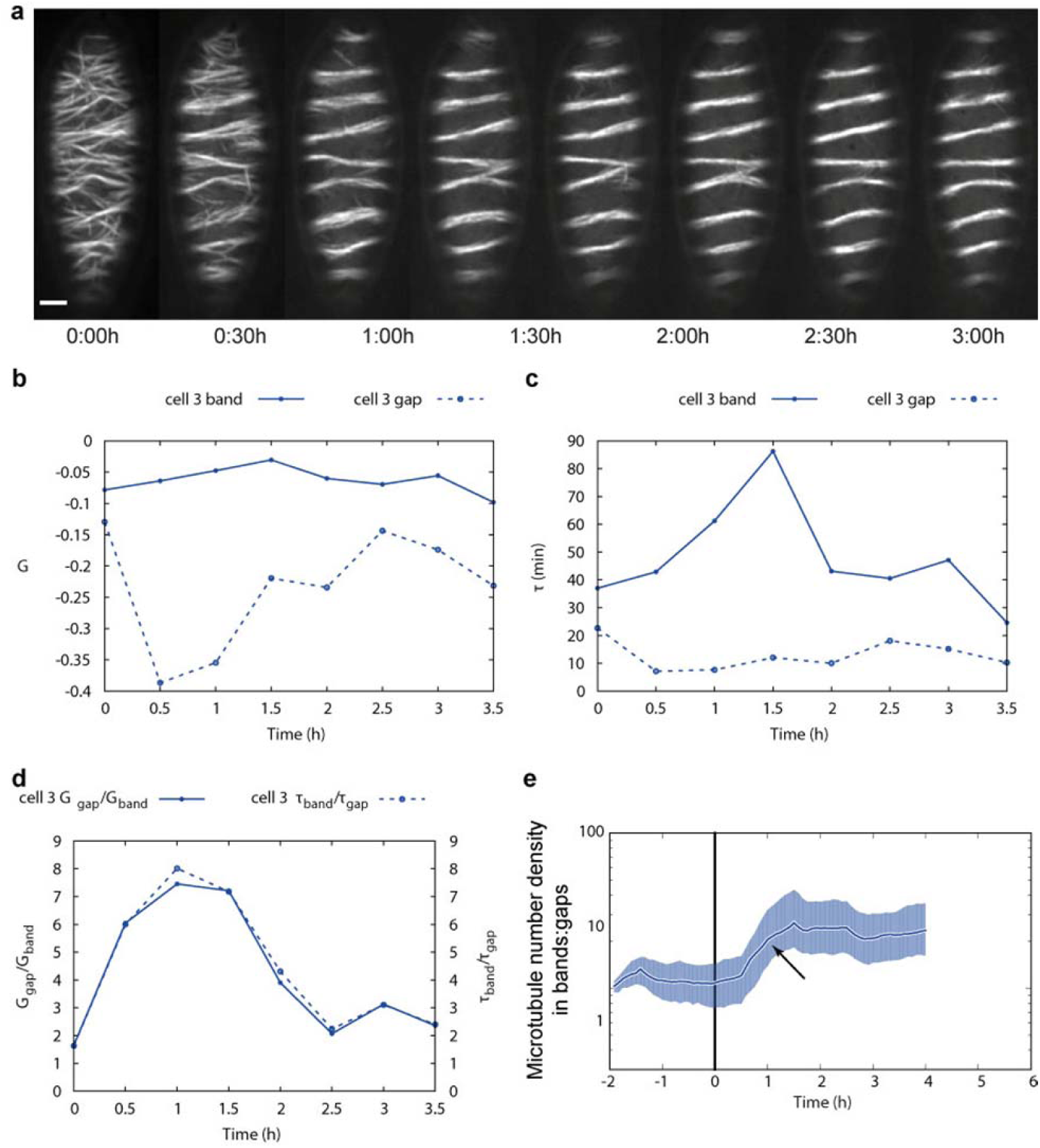
Microtubule re-orientations during the VND7-driven secondary wall program of cell 3. **a**, Average projections of YFP-labelled microtubules in an induced hypocotyl cell. Scale bar = 5 µm. **b-c**, Temporal development of the control parameter *G* (b) and the microtubule lifetime *τ* (c) in bands (solid line) and gaps (dashed line). Microtubules in the bands show more interactions and longer lifetimes than in the gaps. **d**, The temporal development of the ratio of *G*_gap_/*G*_band_ (solid line) and *τ*_band_*/ τ*_gap_ (dashed line) indicates that the number of microtubule-microtubule interactions and the microtubule lifetime needs to be sufficiently different between bands and gaps for separation to occur. **e**, Degree of separation measured as microtubule number density ratio between bands and gaps. Note that a large ratio of *G*_gap_/*G*_band_ and *τ*_band_*/ τ*_gap_ between 0.5 h to 1.5 h in (d) is correlated with a strong increase in the separation rate (arrow). Median ± 16 % and 84 % percentiles. *n* ≥ 100.

**Supplementary Fig. S8.**
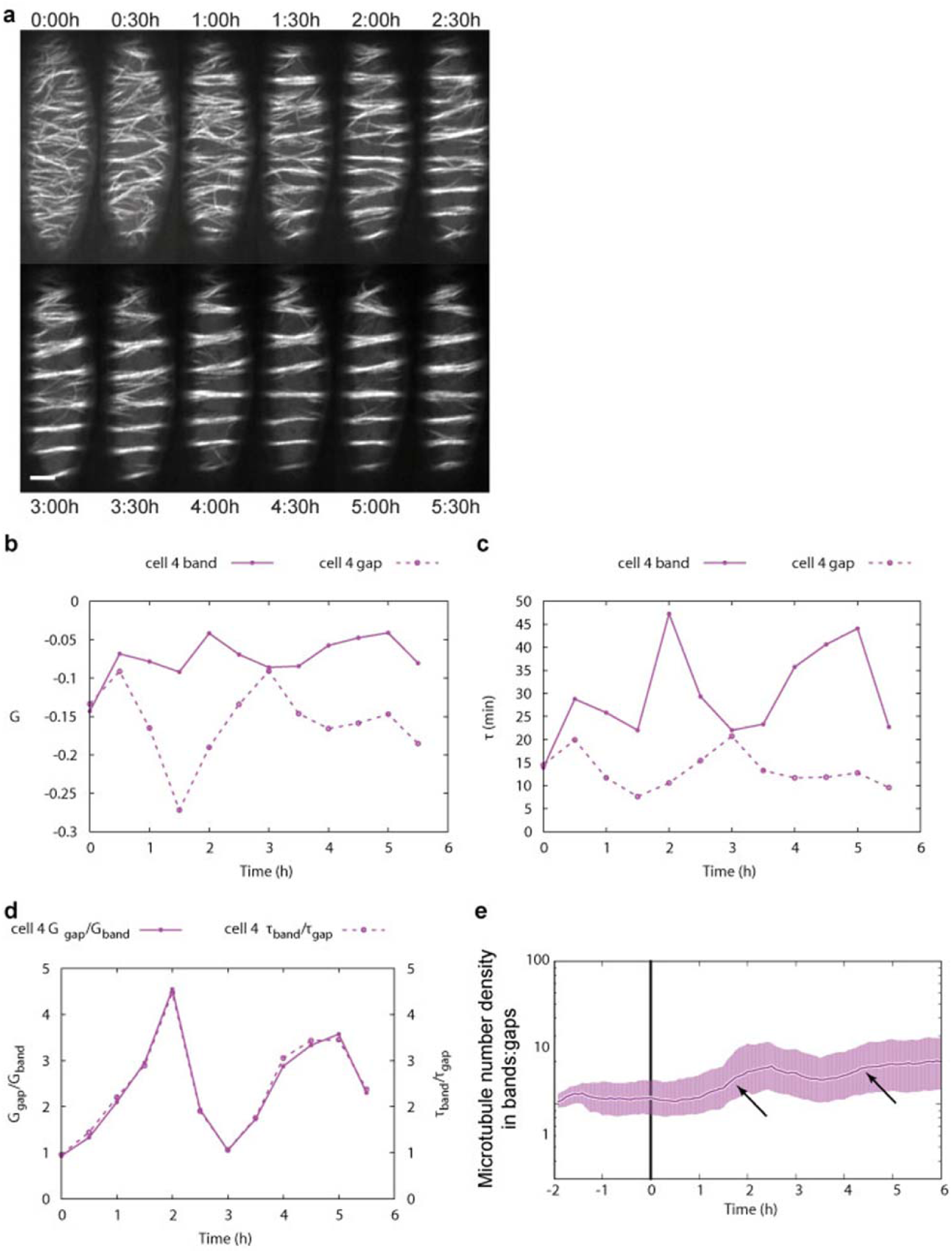
Microtubule re-orientations during the VND7-driven secondary wall program of cell 4. **a**, Average projections of YFP-labelled microtubules in an induced hypocotyl cell. Scale bar = 5 µm. **b-c**, Temporal development of the control parameter *G* (b) and the microtubule lifetime *τ* (c) in bands (solid line) and gaps (dashed line). Microtubules in the bands show more interactions and longer lifetimes than in the gaps. **d**, The temporal development of the ratio of *G*_gap_/*G*_band_ (solid line) and *τ*_band_*/ τ*_gap_ (dashed line) indicates that the number of microtubule-microtubule interactions and the microtubule lifetime needs to be sufficiently different between bands and gaps for separation to occur. **e**, Degree of separation measured as microtubule number density ratio between bands and gaps. Note that a large ratio of *G*_gap_/*G*_band_ and *τ*_band_*/ τ*_gap_ at 2 h and 5 h in (d) is correlated with strong increases in the separation rate (arrows). Median ± 16% and 84% percentiles. n ≥ 100.

**Supplementary Fig. S9.**
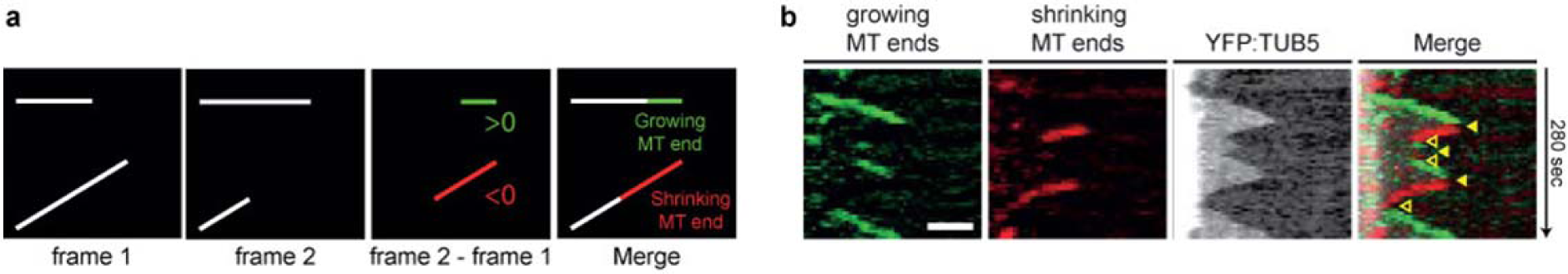
Image subtraction method to detect dynamics of microtubule ends. **a**, Schematic showing the detection of growing (green) and shrinking (red) microtubules (white). A merged image (right panel) allows easy identification of the state, i.e. growing or shrinking, of a microtubule. **b**, The dynamics of microtubules can be visualized using kymograph analysis of a merged image stack containing the growth and shrink information as illustrated in (a), allowing identification of locations where microtubules switched from growing to shrinking (filled arrow heads, termed ‘catastrophe’) and from shrinking to growing (empty arrow heads, termed ‘rescue’). Scale bar = 2 µm.

**Supplementary Fig. S10.**
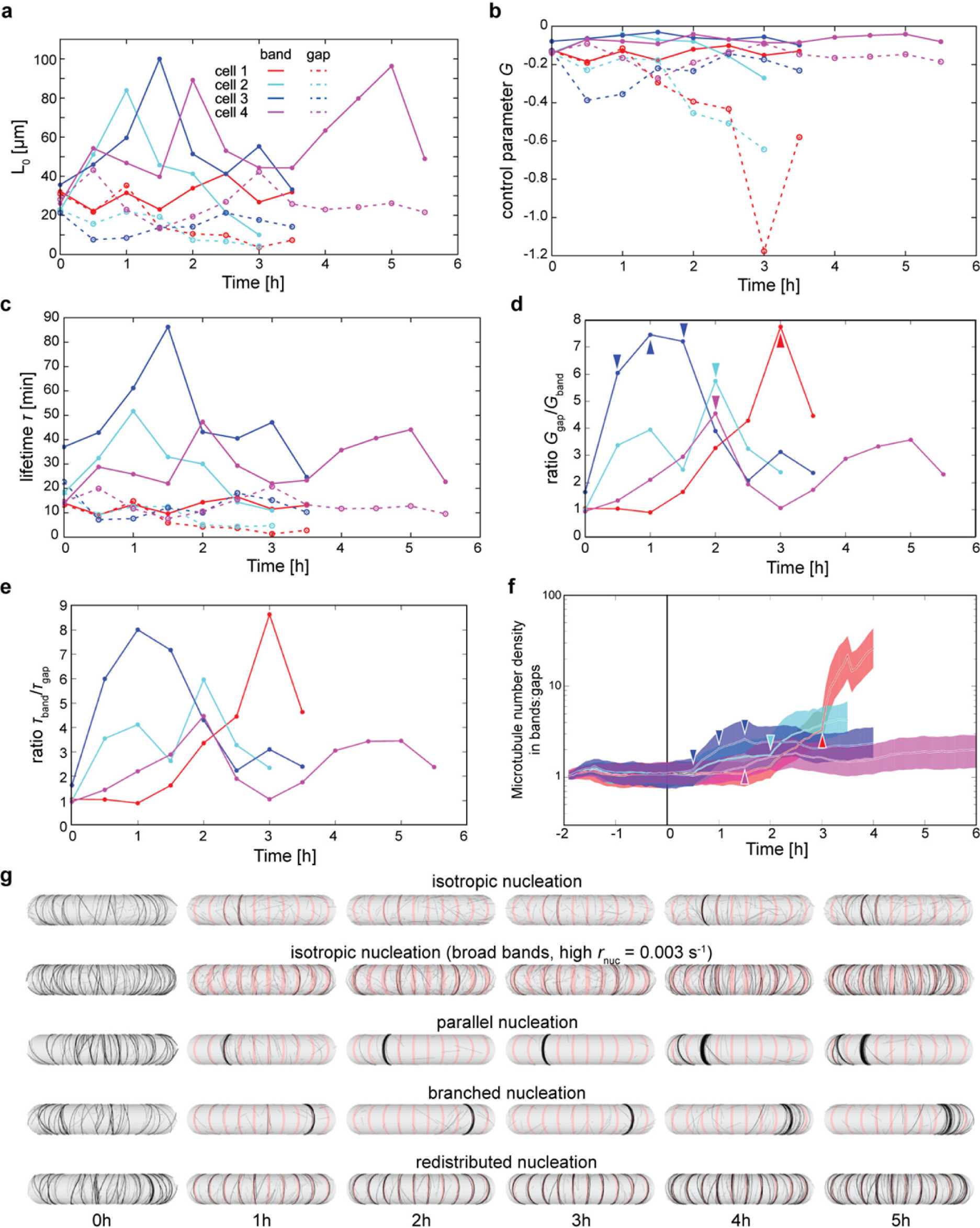
Simulation of microtubule band formation using measured microtubule dynamic instability parameters. **a**, Calculated length of non-interacting microtubules *L*_0_ in band and gap regions for four long-term recordings (see legend for colour-code). During band separation, the calculated average length of microtubules in the bands (solid lines) increases whereas the calculated average length of microtubules in the gaps (dashed lines) is decreased. **b-e**, Average control parameter *G* (b) and average microtubule lifetime *τ* (c) divided into bands (solid) and gaps (dashed), and ratios of *G*_gap_/*G*_band_ (d) and *τ*_band_/*τ*_gap_ (e) over time for the four recorded cells. The peaks in the ratios *G*_gap_/*G*_band_ occur at: cell 1: 3 h, cell 2: 2 h, cell 3: (0.5 h,) 1 h, and 1.5 h, cell 4: 2 h (see coloured arrow heads). Note that the curves in panels a-e start at approximately similar values at T = 0 h indicating that recording began before differences between bands and gaps became dominant. **f**, Simulated degree of separation using the parameters obtained by quantification of the microtubule dynamics in the four analysed cells. Time points of fastest separation: cell 1: 3 h, cell 2: 2 h, cell 3: 0.5 h (and 1 h), and cell 4: 1.5 h (see coloured arrow heads). Median ± 16 % and 84 % percentiles, *n* ≥ 100. **g**, Randomly selected snapshots of representative simulations with the given nucleation mode over a time course of five hours.

**Supplementary Fig. S11.**
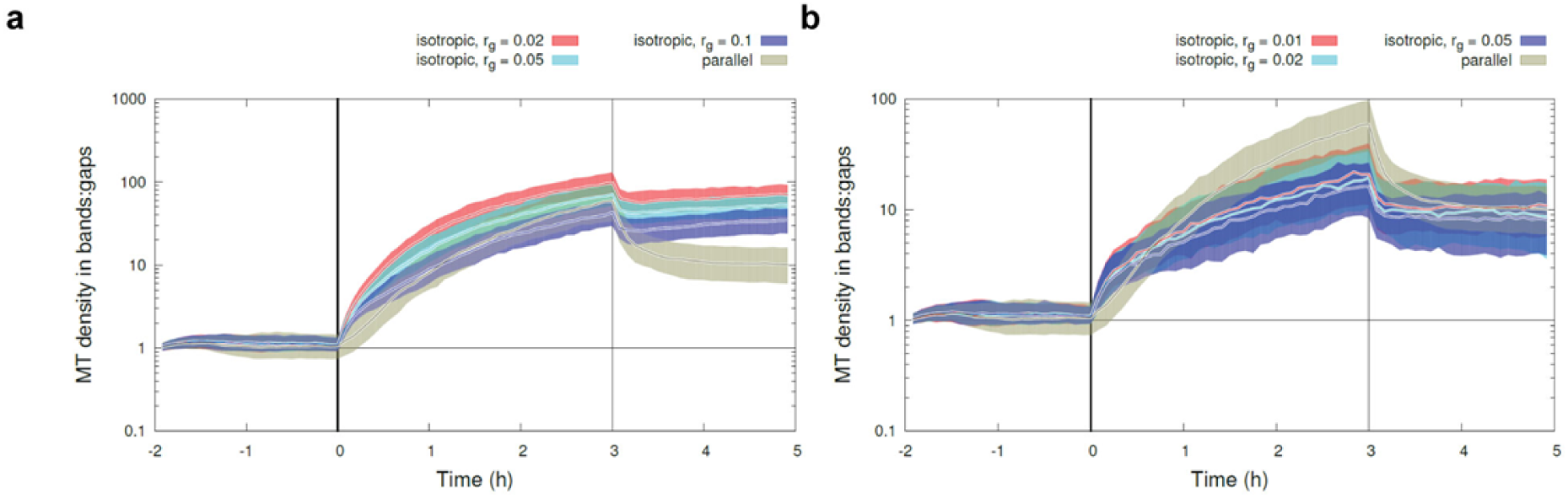
Estimating the feedback strength of microtubule-bound nucleation. **a**, Comparing the default parallel nucleation with isotropic nucleation where rates in gaps were lowered and rates in bands were increased so that *r*_n,gap_ = *r*_g_ × *r*_n,band_. The overall nucleation rate was unaltered. The difference of nucleation rates in bands and gaps was 10-fold (blue, *r*_g_ = 0.10), 20-fold (cyan, *r*_g_ = 0.05), and 50-fold (red, *r*_g_ = 0.02). Isotropic nucleation with a 20-fold rate difference between bands and gaps yields a similar degree of separation as parallel, microtubule-bound nucleation. Median ± 16 % and 84 % percentiles, *n* = 400. **b**, Comparing the default parallel nucleation with isotropic nucleation where only the rates in gaps were lowered and rates in the bands remained constant. The overall nucleation rate was unaltered. The difference of nucleation rates in bands and gaps was 20-fold (blue, *r*_g_ = 0.05), 50-fold (cyan, *r*_g_ = 0.02), and 100-fold (red, *r*_g_ = 0.01). Parallel, microtubule-bound nucleation always yielded better degrees of separation. Median ± 16 % and 84 % percentiles, *n* ≥ 100.

**Supplementary Fig. S12.**
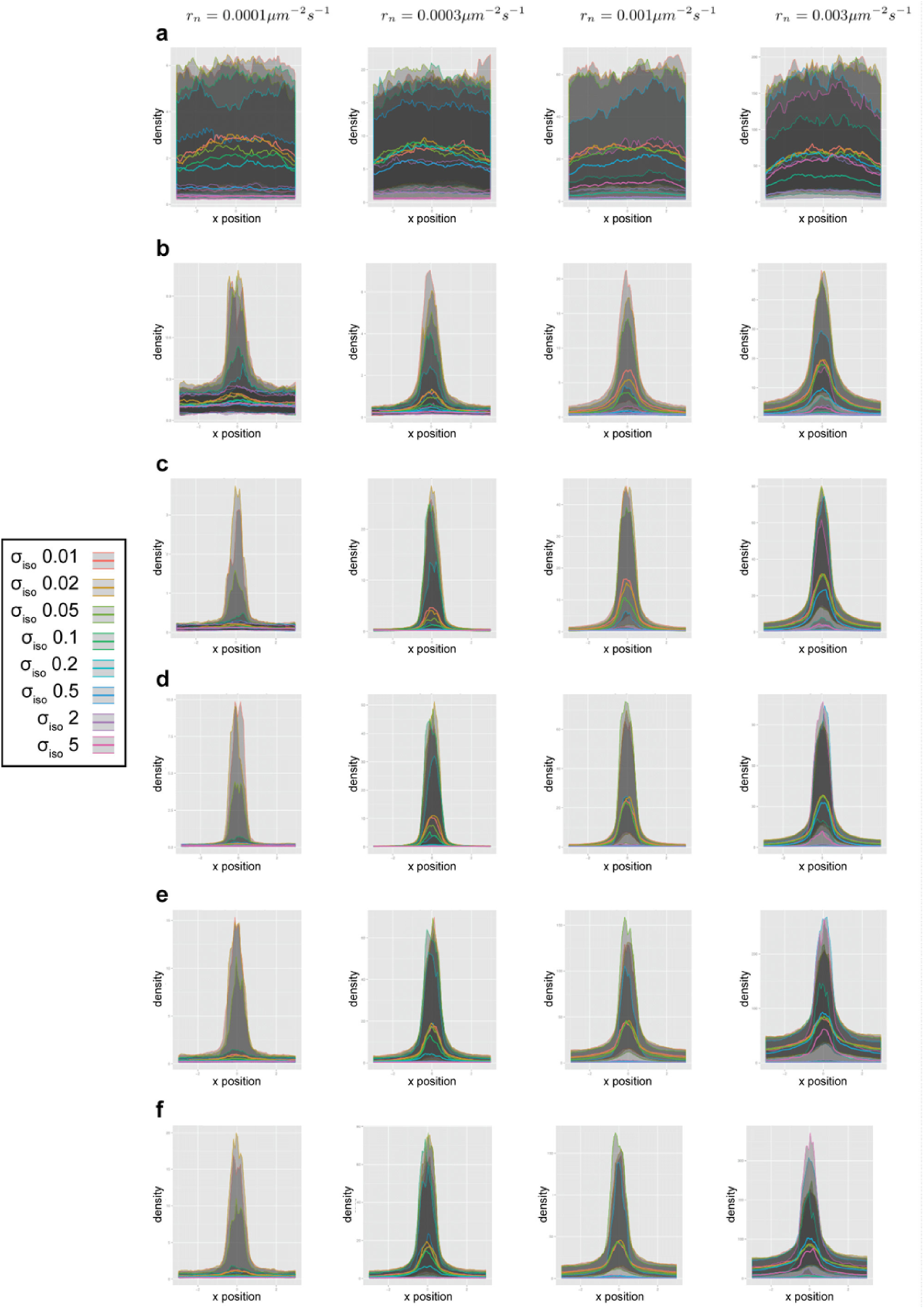
Microtubule density histograms for multiple time stamps, variable *ρ*_iso_ (the density at which 50% of the nucleations occur from microtubules) per nucleation rate *r*_n_. **a**, End of initiation phase (*T* = 0 h); **b**, separation phase (*T* = 1 h); **c**, Separation phase (*T* = 2 h); **d**, Separation phase (*T* = 3 h); **e**, Middle of maintenance phase (*T* = 4 h); **f**, End of maintenance phase (*T* = 5 h). NOTE: similar results were obtained for changing *r*_n_ and treadmilling speed *v*_t_, as both impact total microtubule density.

**Supplementary Figure S13.**
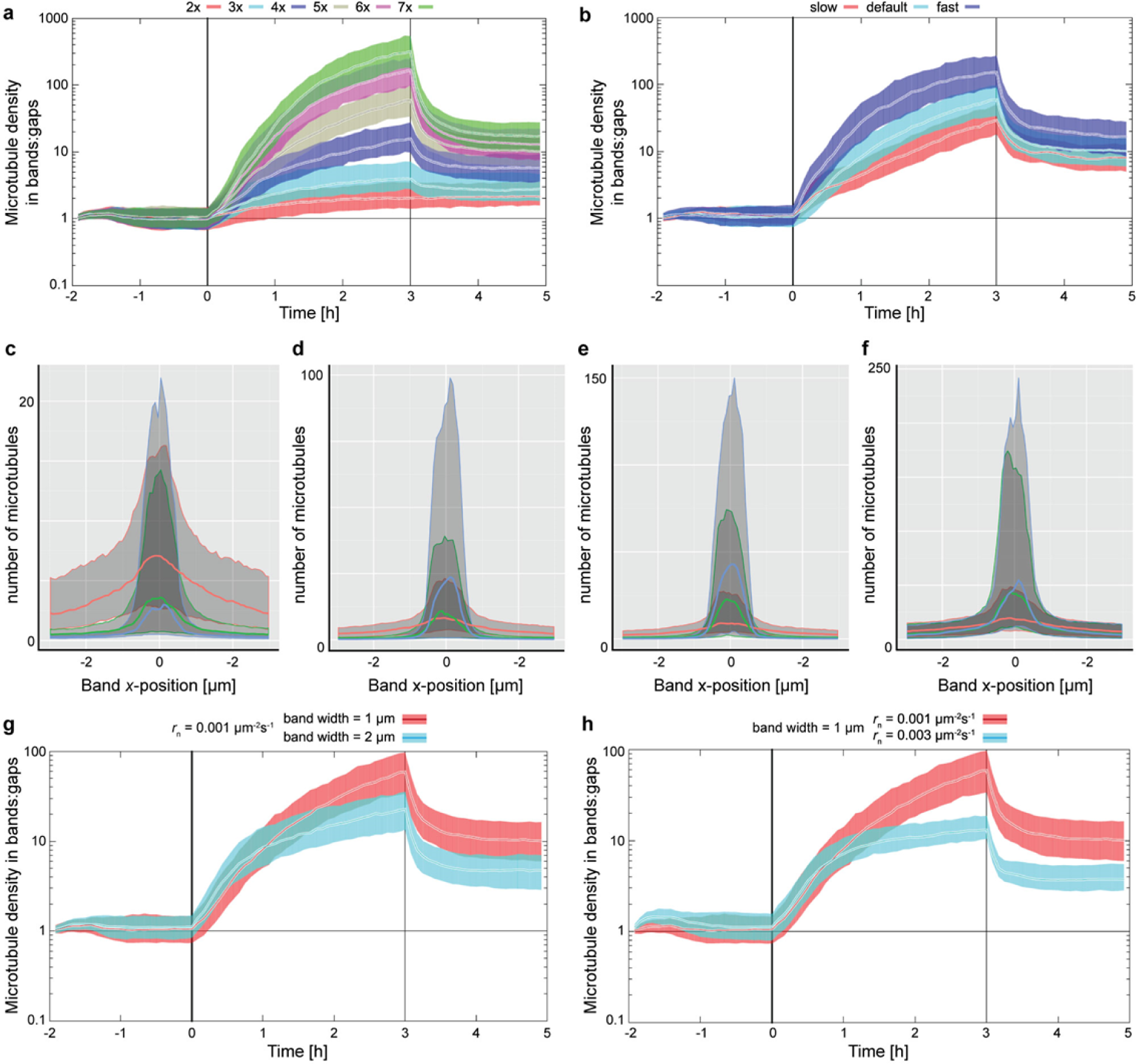
Microtubule stability and turn-over are key determinants of band formation. **a**, Simulated microtubule density ratio between bands and gaps for six different *G*-ratios (separation strengths): two- to seven-fold difference in *r*_cat_ between bands and gaps. Median ± 16% and 84% percentiles, n ≥ 100. **b**, Impact of different microtubule lifetimes *τ* at 5-fold separation strengths. Microtubule lifetimes: 1.9 min (fast), 3.2 min (default), and 5.8 min (slow) in the gaps. Median ± 16 % and 84 % percentiles, *n* ≥ 100. **c-f**, Histograms of microtubule density along the cell axis of a single-banded cell for three different separation strengths (3x red, 5x green, and 7x blue) and four subsequent time points: *T* = 1 h (c), *T* = 2 h (d), *T* = 3 h (e), and *T* = 4 h (f). Note the different scales. Median (thick lines) ± 16 % and 84 % percentiles (thin lines). **g-h**, Increasing the band width (g) from 1 to 2 µm or the nucleation rate (h) from 0.001 to 0.003 µm^−2^ s^−1^ had a negligible impact on the separation rate during the first hour of separation. Median ± 16 % and 84 % percentiles, *n* ≥ 100.

**Supplementary Fig. S14.**
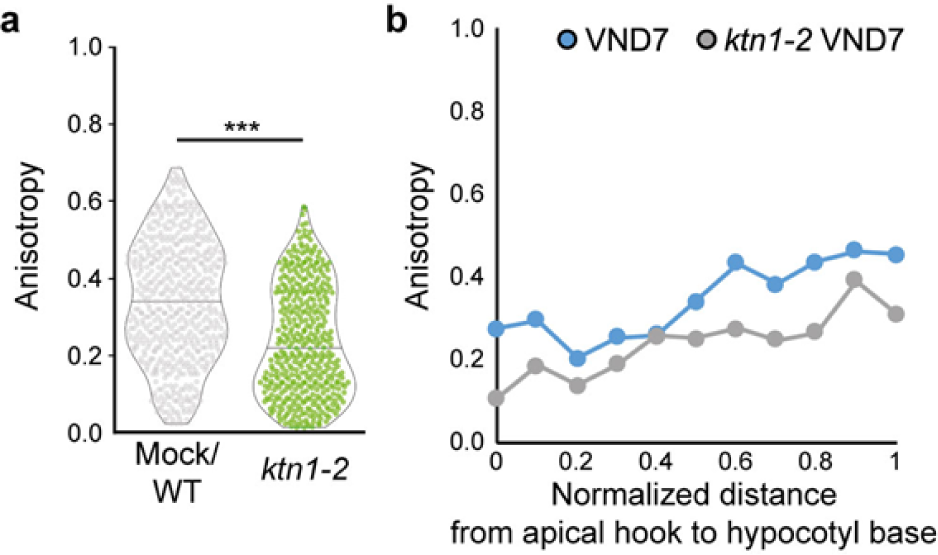
Anisotropy measurements of microtubule arrays in *ktn1-2* versus wild type. **a**, Wild-type seedlings (from Fig. 4e in the main text) showed significantly higher anisotropy (0.34 ± 0.16, mean ± s.d., 563 cells from 7 seedlings) compared to *ktn1-2* (from Fig. 4f in the main text, 0.23 ± 0.14, 422 cells in 6 seedlings) mutants (violin plots indicating median, *p*<0.0001, Welch’s unpaired *t*-test). **b**, Anisotropy of wild-type and *ktn1-2* microtubule arrays normalized to its position along the entire hypocotyl axis from the apical hook (= 0) to the base (= 1). The *ktn1-2* mutant shows consistently reduced anisotropy as compared to wild-type.

**Supplementary Fig. S15.**
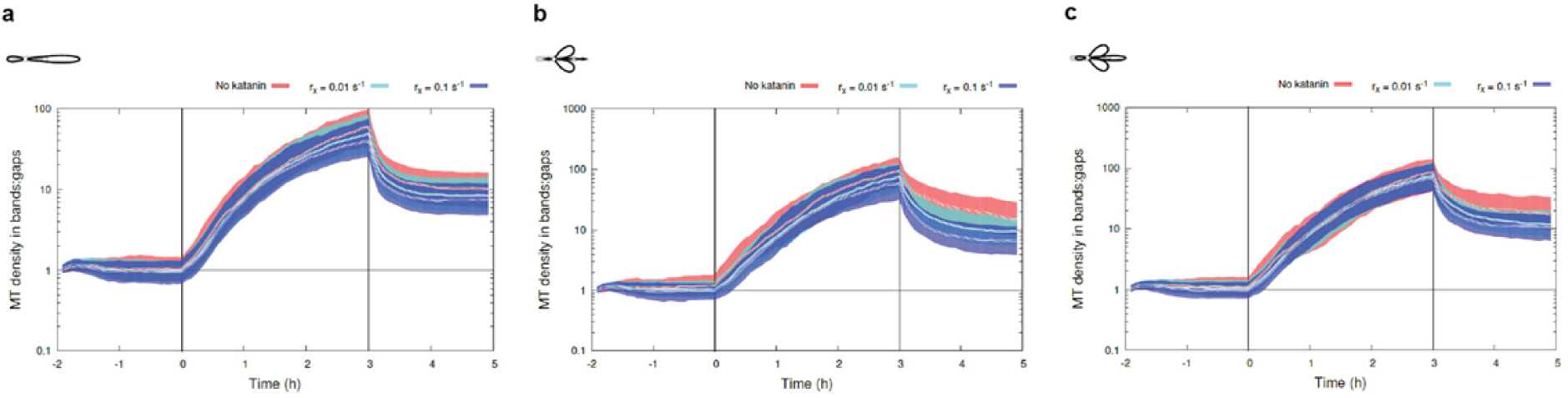
KATANIN activity has little impact on the separation process, independent of the type of nucleation mode. **a-c**, Degree of separation for the parallel (a), branched (b), and fuzzy branched (c; similar to branched, except that small deviations are applied to all microtubule-based nucleations, including the (anti-)parallel ones) nucleation scenario with zero severing rate *r*_x_ = 0 (red line), with low severing rate *r*_x_ = 0.01 s^−1^ (cyan), and with high *r*_x_ = 0.1 s^−1^ (blue). Median ± 16 % and 84 % percentiles, *n* ≥ 100.

**Supplementary Fig. S16.**
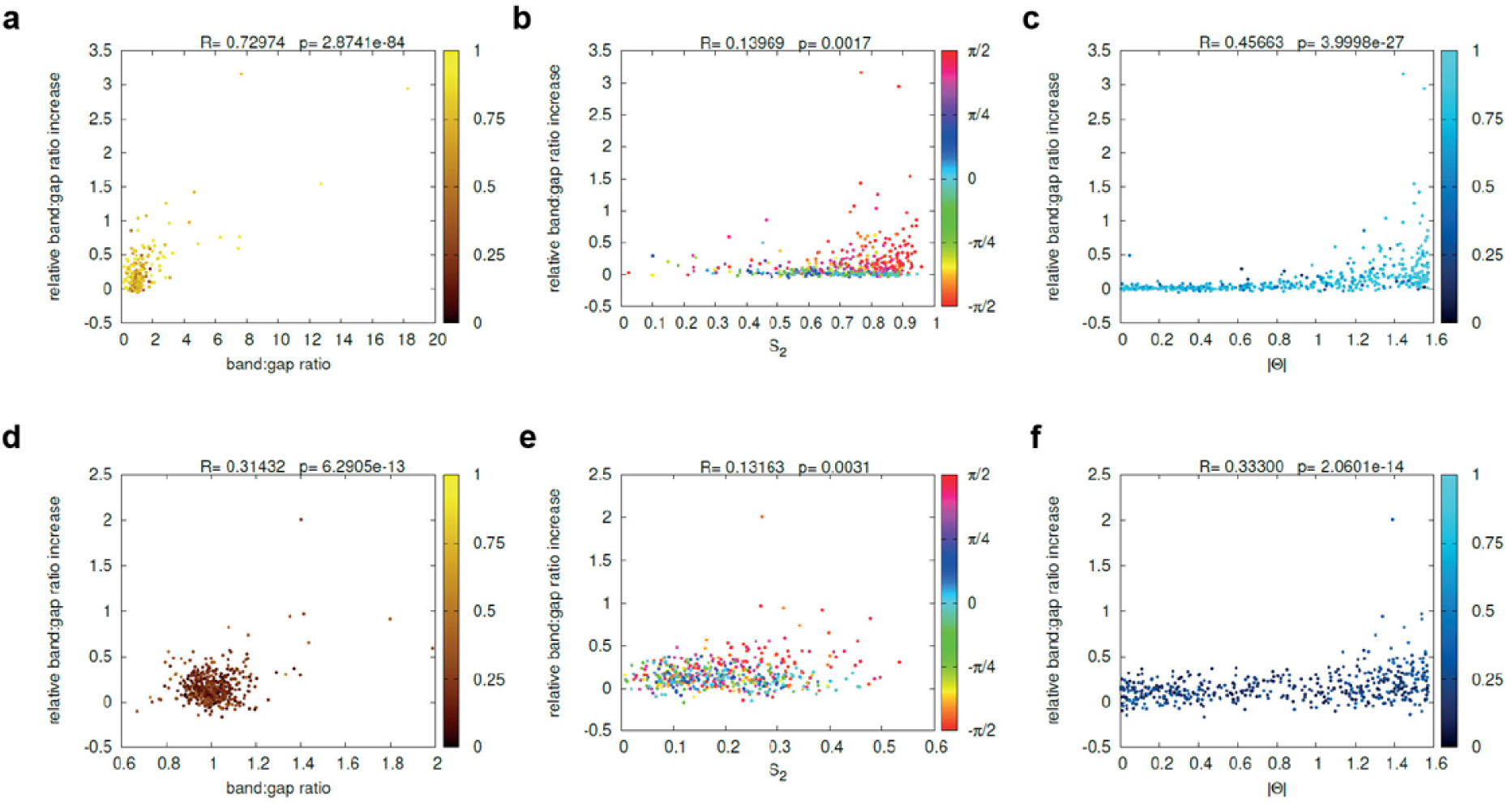
Array orientation and anisotropy impact separation speed *in silico*. **a-f**, Scatter plots of the relative density increase between bands and gaps (= separation rate) and the degree of separation (a,d), the anisotropy parameter S2 (b,e), and the array orientation *θ* (c,f) at *T* = 10 min, for *P*_cat_ = 0.5 (a-c) and for *P*_cat_ = 0.05 (d-f) during initiation phase. Panels a,c,d,f: data points are coloured by the value of the anisotropy parameter S2 indicated. Panels b,e: data points are coloured by *θ* value (in radians) as indicated. Red corresponds to a transverse array.

