## Supplementary Information for "Long-term single-cell imaging and simulations of microtubules reveal driving forces for wall pattering during proto-xylem development"

### Simulation method

All simulations were performed using an extended version of the CorticalSim software (Tindemans and Deinum 2017) (corticalsim v1.26.1; https://zenodo.org/record/801852#.XejjEHt7m5M). In this framework, microtubules are represented as series of connected line segments on the surface of the simulation domain. This two-dimensional surface represents the cell cortex. Microtubules can be in a growing or shrinking state, with stochastic switches between these called rescues and catastrophes. When a growing microtubule impinges on another microtubule, one of three outcomes is possible, depending on the relative angle ('*collision angle*' $\theta$) between both microtubules: (i) continued growth along the obstructing microtubule ('*zippering*' or '*bundling*') for $\theta < \theta^*$ (see Table 1 in this document), (ii) induced catastrophe (with probability $P_{cat}$) or (iii) continued growth in its own direction ('*crossover*') for $\theta > \theta^*$ (Dixit and Cyr 2004; Tindemans et al. 2010).

The simulation domain consisted of a cylindrical cell surface, with radius 7.5 µm and length 60 µm ('*ten-banded cells*') or 6 µm ('*single-banded cells*'). All simulations started with an initiation phase, in which the array was seeded using a transverse orientational bias on the nucleation (30 min), followed by regular nucleation (90 min). During this initiation phase, parameters were the same everywhere on the cell surface. After the initiation phase, the simulation domain was separated into '*band*' and '*gap*' regions with different parameters. The end caps of the cylinders were treated the same as gap regions. Bands were 1 µm wide, with 5 µm wide gaps in-between (ten-banded cells), or a single 1 µm wide band in the centre of the cell (single-banded cells).

For computational efficiency, the number of parameters that differed between bands and gaps was minimized, while maintaining the same values of control parameter G (see section below) in both bands and gaps. When rates differed between regions (spontaneous catastrophe rate $r_{cat}$, and in some simulations also nucleation rate $r_n$), all events were scheduled by the highest rate applicable and proportionally discarded (not executed) in the regions with the lower rate.

With this setup, we performed two kinds of simulations: (i) direct simulations of experimentally observed parameters, where the simulation was subdivided into 30-minute intervals, and caricature simulations, with a simpler structure of a three-hour separation phase (large difference in *G* between bands and gaps) and a two-hour maintenance phase (smaller difference).

### Control parameter G

The control parameter *G* is a theory-derived quantity (Tindemans et al. 2014; Tindemans et al. 2010) that combines the microtubule dynamic instability parameters and nucleation rate into a single number:

$$G = \sqrt[3]{\frac{2(v^+ - v^t)^2\,(v^- + v^t)}{r_n v^+ (v^+ + v^-)}} \times \left(\frac{r_{res}}{v^- + v^t} - \frac{r_{cat}}{v^+ - v^t}\right) \qquad (1)$$

with $v^+$ the plus-end growth velocity, $v^-$ the plus-end shrinkage velocity, $v^t$ the minus-end retraction velocity, $r_{res}$ the rescue rate, $r_{cat}$ the spontaneous catastrophe rate and $r_n$ the nucleation rate. *G* consists of the negative ratio of two length scales: the '*interaction distance*' or average distance between consecutive encounters of growing microtubule tips (first term in Eq. 1) and the '*average microtubule length*' in the absence of microtubule-microtubule interactions ($L_0$; second term). The closer *G* approaches zero, the more interactions will take place during an average microtubule lifetime and hence, the larger the propensity of the array to align spontaneously (Tindemans et al. 2014; Tindemans et al. 2010; Hawkins et al. 2010). Typically, spontaneous alignment will occur if $G > G^*$, with $G^*$ being the critical *G*-value that depends on the specifics of the microtubule interactions and angle-dependent collision outcomes. If $G > 0$, the system is in a regime of unbounded microtubule growth. Consequently, a biological microtubule array must always converge to a state where $G < 0$, although *G* may be positive during the initial stages of array establishment (Tindemans et al. 2014). For all parameter combinations in our simulations, experimentally observed parameters naturally satisfy $G < 0$. Several simulation studies showed that *G* constitutes a powerful concept in understanding array behaviour (Deinum et al. 2011; Tindemans and Deinum 2017; Deinum and Mulder 2013).

### Array alignment and orientation: order parameter S2

The global degree of alignment of the array can be measured using nematic order parameter S2 computed on the body of the cylindrical simulation domain.

$$S_2 = \sqrt{\langle \cos(2\theta) \rangle^2 + \langle \sin(2\theta) \rangle^2} \tag{2}$$

with $\langle \ldots \rangle$ denoting a length-weighted ensemble average and $\theta$ the orientation angle of individual microtubules. The overall orientation of the array can then be computed as:

$$\theta = \tan^{-1}\left(\frac{\langle \sin(2\theta) \rangle}{\langle \cos(2\theta) \rangle + S_2}\right) \tag{3}$$

**Constant simulation parameters**

Simulation parameters that remain constant during individual simulations are given in Table 1 in this document.

<u>Nucleation modes:</u>

**Isotropic** Nucleation uniformly distributed over the simulation domain, with uniformly distributed nucleation angles.

**Parallel (default)** A fraction of the nucleations, $\frac{\rho}{\rho_{iso}+\rho}$, with $\rho$ the average microtubule density and parameter $\rho_{iso}$ the density at which 50% of the nucleations occur from microtubules, is nucleated from existing microtubules, with location proportional to microtubule density, and the same orientation of the parent microtubule (93% parallel, 7% anti-parallel) ± a small deviation following the relative surface area of an ellipse with eccentricity 0.89. For further details of this nucleation mode see (Deinum et al. 2011)

**Branched** Similar to parallel nucleation, but 62% is nucleated with an average angle of 35 degrees relative to the parent microtubule, with a small deviation. The remaining parallel (31% parallel, 7% anti-parallel) component is nucleated exactly along the parent microtubule. This mode is designed to best resemble the experimental data by Chan et al., (2009) (Chan et al. 2009; Deinum et al. 2011).

**Fuzzy Branched** Similar to branched, except that small deviations are applied to all microtubule-based nucleations, including the (anti-)parallel ones.

**Redistributed** Nucleation position and angle are determined as they are in parallel nucleation, followed by a shift of an integer times the width of a band + gap. Consequently, distance to the centre of the nearest band is conserved, but global position is redistributed. This nucleation mode is used solely to demonstrate the need to break the global competition for nucleation sites among bands in order to obtain a pattern with ten-banded microtubule cells.

**Table 1: Constant simulation parameters**

| Parameter | Value | Description | Reference |
|---|---|---|---|
| $v^t$ | 0.01 µm/s | Minus-end retraction velocity | (Shaw et al. 2003; Deinum et al. 2011) |
| $r_x$ | 0 $s^{-1}$ | Severing rate at microtubule crossovers, per crossover. Other used values: 0.1, 0.01 $s^{-1}$. | (Deinum et al. 2017) |
| $r_n$ | 0.001 $\mu m^{-2}s^{-1}$ | Nucleation rate, average per unit surface area. | (Tindemans et al. 2014) |
| $\Theta^*$ | 40° | Critical angle for zippering/bundling: after collisions with relative angles of $\theta < \theta^*$, the impinging microtubule continues it growth along the obstructing microtubule. | (Dixit and Cyr 2004) |
| $P_{cat}$ | 0.5 | Fraction of collisions with relative angle $\theta > \theta^*$, resulting in induced catastrophe. | (Deinum et al. 2011) |

### Determination of band/gap-specific parameters

<u>Cell simulations</u>

Parameters $v^+$, $v^-$, $r_{cat}$, and $r_{res}$ were based on experimentally measured parameters from this study. For the simulation of the individual cells, the averages measured in bands were used as is. For the gaps, the band values were used for parameters $v^+$, $v^-$, and $r_{res}$, while $r_{cat}$ was chosen such that the calculated $G$-value (using $v^t$ = 0.01 µm/s everywhere) remained the same, i.e., solving $r_{cat,gap,adjusted}$ from:

$$\sqrt[3]{\frac{2(v_{gap}^{+}-v^{t})^{2}\,(v_{gap}^{-}+v^{t})}{r_{n}v_{gap}^{+}(v_{gap}^{+}+v_{gap}^{-})}}\times\left(\frac{r_{res,gap}}{v_{gap}^{-}+v^{t}}-\frac{r_{cat,gap}}{v_{gap}^{+}-v^{t}}\right)$$
$$=\sqrt[3]{\frac{2(v_{band}^{+}-v^{t})^{2}\,(v_{band}^{-}+v^{t})}{r_{n}v_{band}^{+}(v_{band}^{+}+v_{band}^{-})}}\times\left(\frac{r_{res,band}}{v_{band}^{-}+v^{t}}-\frac{r_{cat,band}}{v_{band}^{+}-v^{t}}\right) \quad (4)$$

with subscripts *band* or *gap* indicating the experimentally measured values in the respective regions (see Supporting Table S1-S4 for the resulting values). Note that nucleation rate $r_n$ cancels out of this equation, hence the same $r_{cat,gap,adjusted}$ holds for every $r_n$. We have chosen this approach over directly using all measured parameters, because it keeps the internal structure of the simulations much simpler, saving extensive bookkeeping, complex programming and computation time.

<u>Caricature simulations</u>

The baseline parameters for the caricature were roughly based on the average values of the experimental cells. Although in some cells the *G*-values in bands increased during the separation process, we kept them constant at a level representative of early arrays. From this baseline, the $r_{cat}$ value in gaps was increased by an integer factor (default 5x during separation phase and 2x during maintenance phase). The resulting differences in *G*-value were similar to the large differences observed in the experimental cells.

**Statistical analysis of simulation results**

Analysis was performed using custom written python and bash scripts.

<u>Ten-banded cells</u>

Per parameter set, at least 100 independent cells were simulated. The microtubule density along the cell axis was tracked during the simulation in histograms with 120 bins (i.e., 0.5 µm wide), recorded every five minutes. On these histograms, the degree of separation was calculated as the ratio of average microtubule density in the band region: gap region, where the 1 µm wide regions flanking the bands (‘*edge regions*’) were excluded from the analysis. On this statistic, we calculated median and

16th and 84th percentiles, the latter two corresponding to ±1$\sigma$ for normally distributed data.

Single-banded cells

Per parameter set, at least 2000 single-banded cells were simulated. Of these, the cells with longitudinal orientation were discarded ($|\theta| < 0.6$, with $\theta$ the array orientation angle). This was necessary for the single-banded cells only, because these were far more prone to lose, or not establish a transverse orientation compared to the ten-banded cells. Discarded simulations were never more than 50% of the total). The remaining cells were randomly distributed into groups of ten, thus simulating a ten-banded cell with ten independent bands. (Up to 9 single-banded cells were discarded in this process.) This resulted in 100-200 independent composite cells. Degree of separation and percentiles were computed as if these composites were regular ten-banded cells. Density histograms had 120 (0.05 µm wide) bins per single-banded cell, so 1200 per composite cell.

Density histograms (single-banded cells)

Averaged density histograms were computed based on at least 400 independent simulations per parameter set. Simulations with a longitudinal orientation were discarded (see above). On the remaining data, the median and the 16th and 84th percentiles were determined per bin of the spatially resolved density histograms. For improved visualization, the curves were computed based on the data for percentile ±2 (e.g., 14th - 18th for the 16th.) followed by smoothing with a 91th order spline. Spline order was chosen so high to keep most of the noise characteristics of the data but obtain slightly smoother-looking curves. This analysis was implemented in R statistical language.

**Quantification of the strength of the nucleation feedback**

To quantify the strength of the positive feedback we compared parallel microtubule-bound nucleation to microtubule-independent isotropic nucleation with different nucleation rates in bands and gaps. We decreased the nucleation rate in the gaps and increased it in the bands, such that the average nucleation rate remained the same. When we decreased the nucleation rate in the gaps, we found that parallel nucleation initially showed delayed separation as compared to isotropic nucleation but subsequently reached a similar degree of separation after three hours (Supplementary

Fig. S11). This indicates that an approx. 10- to 20-fold difference in the nucleation rate between bands and gaps would be needed for isotropic nucleation to reach a similar degree of separation as in the case of microtubule-bound parallel nucleation. However, when nucleation rates were lowered only in the gaps while keeping the nucleation rates in bands constant, we observed a smaller degree of separation (Supplementary Fig. S11). This finding indicates that the increase of nucleation in bands contributes to a fast separation process. Given this strong feedback, one would expect that the affinity of nucleation complexes for microtubules would have a strong impact on the degree of separation reached in the simulations. This, however, was not the case: within a large range of the microtubule affinity parameter $\rho_{\text{iso}}$ (the global density at which 50% of the nucleations occurs from existing microtubules), the final degree of band separation hardly depended on this affinity (Supplementary Fig. S12), where the suitable bandwidth strongly depended on the nucleation rate (which has a large impact on microtubule density). Notably, in those cases the median band densities always ended many times higher than $\rho_{\text{iso}}$, suggesting that as time progresses, the number of nucleation complexes rather than their affinity for microtubules determines the amount of nucleation in the bands.

**Simulations identify two major factors that affect microtubule separation rates**

We next set out to assess whether our simulation platform could aid in determining factors that impacted the microtubule separation rate during trans-differentiation. The first obvious parameters to vary were the ratios of *G* and *τ*. To this end, we varied the ratio of $r_{cat,gap}$ to $r_{\text{cat,band}}$ with values between two to seven during the separation phase). We found that the corresponding ratio $G_{\text{gap}}/G_{\text{band}}$ – the '*separation strength*' – substantially impact both the initial separation rate and the total degree of separation, which peaked at 3 hours (Supplementary Fig. S13). In an attempt to decouple *G* and *τ*, we next computed new caricature parameter sets that have the same *G* and *τ* ratios during the separation phase (at a default 5x separation strength), but with different microtubule life-times. We found that a faster turnover time results in a faster separation, even if the *G* and *τ* difference between bands and gaps is constant (Supplementary Fig. S13). We also tested additional factors and found that the separation rate was largely independent of the band width and overall nucleation rate during the first hour of separation (Supplementary Fig. S13).

**Microtubule depolymerization in bands is typically the limiting factor of separation rate**

The above simulation results indicate that the values and/or ratios of both $G$ and $\tau$ are important determinants of the separation process. We therefore returned to the four full-length experimental time series. For computational tractability, we converted each of the measured data sets to simulation parameters by calculating the $r_{cat,gap}$ that maintains the same $G$-value in the gaps while using the other measured parameters from the bands (Supplementary Tables S1-S5). We then simulated all four cells and compared the separation process with notable characteristics in the temporal profiles of $G$, $\tau$ and the ratios of $G_{gap}/G_{band}$ and $\tau_{band}/\tau_{gap}$ of these cells (Supplementary Fig. S10 and S5-S8). Notably, the ratios $G_{gap}/G_{band}$ and $\tau_{band}/\tau_{gap}$ followed very similar profiles for all cells.

We observed that the time points of fastest separation and the time points of highest separation strength are closely related in cell 1 and cell 2 (see arrow heads in Supplementary Fig. S10). In cell 3 and 4, the fastest separation occurs earlier and coincides with the periods of lowest $G_{gap}$ and $\tau_{gap}$, which is also the case for cell 1. This suggests that the breakdown of microtubules in the gap regions is the limiting factor of the separation process. Moreover, cell 4 overall has the most stable microtubules in the gaps (highest $G_{gap}$ and $\tau_{gap}$) and shows the slowest and overall least degree of separation. Only for cell 2, the fastest separation occurs before the lowest $G_{gap}$. There are two, not mutually exclusive, explanations for this: 1) the differences in $G$ at 2h (peak separation rate) and the two following time slots are not that large (-0.31, -0.35 and -0.45, respectively) and differences in $\tau_{gap}$ are even smaller (5.0, 4.4, and 4.7 minutes, respectively) so saturation of the separation process may play a role. Notably, the separation already levels off during the period of 2 to 2.5h. 2) During the last two time points (2.5 and 3h), the $G_{band}$ values of cell 2 decrease to the lowest level observed in bands in this study. This could indicate that stability in the bands hardly affects the separation process, so long as it is sufficiently high, and that these final levels in cell 2 no longer are sufficiently high. Overall, we can conclude that the rate-limiting factor of the separation process in these simulated experimental cells is the breakdown of microtubules in the gaps.

**Example parameter file (default parameters, caricature, separation phase)**

CorticalSim parameters:

```
corticalSim 1.32
outputDir
/data/microtubules/bands/catBands/separation/singleBand/charic
ature2/rhoIso/data/gridcylin
newParameterReadInterval 10800
newParameterFile params-3.txt
vPlus 0.05
vMin -0.08
vTM 0.01
kSev 0
kCross 0
kCat 0.0016 0.008
kCatOrient x
bandGapWidth 1 5
nSpirals 0
catPattern singleBand
spiralPitch 0
wrapLength 1
projectedPeriod 6
projectedBand 1
calcNucBiasAngle 0
redistributeNucleationOverBands 0
redistributeShift 0
redistributeShiftNumber 0
kRes 0.001
poolDensity 10
treadmilling 1
severing 0
crossSevering 0
catastropheMultiplier 1
forbiddenZones 0
```

```
restrictedPool 0
edgeCatastropheEnabled 0
edgeCatastropheSmooth 0
pCatRegularEdge 0
pCatSpecialEdge 0
kNuc 0.001
nucleationType ellipse
nucleationBiasType ellipseApolar
nucleationAlpha 0
nucleationBiasAngle 1.5708
zippering 1
ind_cat 1
proportionalCatastrophes 0
induced_cat_fraction 0.5
cat_start_angle 40
zipFraction 1
magic_angle 40
c0Value 0.75
z0Value 1
interactionType zipFirst
bundleType simple
discreteAngleNumber 0
random_seed 1142209023
stopTime 25199
measurementInterval 300
densityLimit 1e+101
wallClockLimit 1e+10
memoryLimit 450
movieEnabled 0
angleHistogramBins 20
angleHistogramOptical 1
hiresLengthHistogramBins 0
loresLengthHistogramBins 0
loresAngleHistogramBins 0
loresLifetimeHistogramBins 0
```

hiresLifetimeHistogramBins 0

histogramAverageSamples 0

spatialHistogramType x

spatialHistogramBinsX 120

spatialHistogramBinsY 45

spatialHistogramCountCaps 0

spatialHistogramWriteRaw 0

geometry gridcylinder 6 7.5 2

ellipseReducedFreeRate 0

ellipseReducedFreeRateAcceptFraction 1

ellipseEpsilon 0.89

ellipseForwardAlongmicrotubule 0

ellipseLeftFraction 0.31

ellipseRightFraction 0.31

ellipseBackwardFraction 0.07

ellipseSidewaysAngle 0

nucleationHalfIsotropicDensity 0.5

Calculated theory parameters:

c0 0.487679

z0 0.297881

x0 0.487679 (should be: 0.487679)

G_0 -0.110119

adjustedG -0.102225

l0 3.8118

Gmax -0.102225

kCatMin 0.0016

Gmin -0.668394

kCatMax 0.008
